# Translational profiling of mouse dopaminoceptive neurons reveals a role of PGE2 in dorsal striatum

**DOI:** 10.1101/2020.09.02.279240

**Authors:** Enrica Montalban, Albert Giralt, Lieng Taing, Yuki Nakamura, Claire Martin, Benoit de Pins, Assunta Pelosi, Laurence Goutebroze, Laia Castell, Wei Wang, Kathrina Daila Neiburga, Letizia Vestito, Angus C. Nairn, Emmanuel Valjent, Denis Hervé, Nathaniel Heintz, Nicolas Gambardella Le Novère, Paul Greengard, Jean-Pierre Roussarie, Jean-Antoine Girault

## Abstract

Forebrain dopaminoceptive neurons play a key role in movement, action selection, motivation, and working memory. Their activity is dysregulated in addiction, Parkinson’s disease and other conditions. To characterize the diverse dopamine target neuronal populations, we compare translating mRNAs in neurons of dorsal striatum and nucleus accumbens expressing D1 or D2 dopamine receptor and prefrontal cortex expressing D1 receptor. We identify D1/D2 and striatal dorso-ventral differences in the translational and splicing landscapes, which establish the characteristics of dopaminoceptive neurons. Expression differences and network analyses identify novel transcription factors with presumptive roles in these differences. Prostaglandin E2 appears as a candidate upstream regulator in the dorsal striatum, a hypothesis supported by converging functional evidence indicating its role in enhancing D2 dopamine receptor action. Our study provides powerful resources for characterizing dopamine target neurons, new information about striatal gene expression patterns, and reveals the unforeseen role of prostaglandin E2 in the dorsal striatum.

## INTRODUCTION

In multicellular organisms, differentiation results from the acquisition of different patterns of gene expression by each cell type. Identifying the mRNA that are translated in a cell population provides clues about its functional properties as well as its vulnerability to pathological conditions. Knowledge of selectively expressed genes also offers a means to target a cell population for investigative or therapeutic purposes. The nervous system represents a major challenge in terms of cell diversity and the definition and number of different cell types is still an open question^1^. Dopamine (DA) exerts neuromodulatory effects on large brain regions containing many cell types, including the striatum and the prefrontal cortex (PFC)^2^. The importance of DA in the physiology of motor circuits, reward process, and working memory, as well as in a wide variety of pathological conditions is thoroughly documented^3^. DA reduction results in Parkinsonian syndromes, whereas its repeated increase by drugs of abuse is a key element leading to addiction^4,5^. DA is also involved in many other conditions ranging from hyperactivity and attention deficit disorder to schizophrenia.

Among the 5 types of DA receptors, the D1 and D2 receptors (DRD1 and DRD2) are the most abundant in the striatum and are also expressed, at much lower levels, in the PFC^6^. In the PFC, DRD1 and DRD2 are found in pyramidal cells as well as in GABAergic interneurons^7,8^. In the rodent striatum, DRD1 and DRD2 are mostly expressed in striatal projection neurons (D1- and D2-SPNs, a.k.a. medium-size spiny neurons, MSNs), which have different functional properties but work in an integrated manner to shape behavior^9^. In the dorsal striatum (DS), D1-SPNs innervate the substantia nigra and the internal (medial) globus pallidus and form the direct pathway, whereas D2-SPNs innervate the external (lateral) globus pallidus providing the first link in the indirect pathway^10^. In the nucleus accumbens (NAc) receptor expression pattern and neuronal connections are less dichotomic^11,12^. DRD2 is also expressed, at lower levels, in cholinergic interneurons^13^. D2-SPNs are the first altered in Huntington’s disease^14^ and DRD2 are decreased in chronic addiction^15^. DS and NAc have different functions and roles in pathology^16^. In contrast to the global differences in gene expression between D1- and D2-SPNs, first shown for substance P and enkephalin^10^, which are well documented^17–20^, little is known about differences between dorsal and ventral D1- and D2-SPNs. Single cell RNA sequencing emphasized the existence of multiple striatal cell populations^21–23^ but did not provide in-depth characterization of regional differences, and PFC neurons expressing DRD1 have not been investigated.

To address regional differences in dopaminoceptive cells, we use translating ribosome affinity purification (TRAP) in transgenic mice expressing enhanced green fluorescent protein (EGFP) fused to L10a ribosomal protein (Rpl10a)^24,25^ under the control of the *Drd1* or *Drd2* promoter^18^. Polysomes from the cell population in which the promoter is active are isolated by EGFP-immunoprecipitation providing access to actively translated mRNAs, also known as the “translatome”^18,24,25^. Here, we combine TRAP with RNAseq (TRAP-Seq) to characterize dopaminoceptive neurons in depth. We find major differences in mRNA expression and isoform/splicing profiles in PFC, DS, and NAc, including similarities between D1 and D2 populations depending on their striatal localization. This work provides a comprehensive data set that can be used to identify the expression pattern of any gene of interest in dopaminoceptive cells. Network analysis identifies known and novel transcription factors potentially involved in striatal regional specification. Analysis of upstream regulators points to the potential role of prostaglandin E2 (PGE2) in the DS and we provide evidence for an important modulatory role of this lipid mediator.

## RESULTS

### Data quality

Drd1-EGFP/Rpl10a and Drd2-EGFP/Rpl10a TRAP mice expressed high levels of EGFP/L10a in the striatum (**Fig. 1a**) with a pattern consistent with the previously described expression in D1- and D2-SPNs^18,26^. In the PFC only Drd1-EGFP/Rpl10a mice expressed sufficient amounts of EGFP-Rpl10a to allow polysome purification. We studied by TRAP-Seq polysome-associated mRNA from the PFC, DS, and NAc of Drd1-EGFP/Rpl10a mice and the DS and NAc of Drd2-EGFP/Rpl10a mice (**Fig. 1b**), using 14-19 independent samples per population (**Supplementary Table 1a**).

**Fig. 1:**
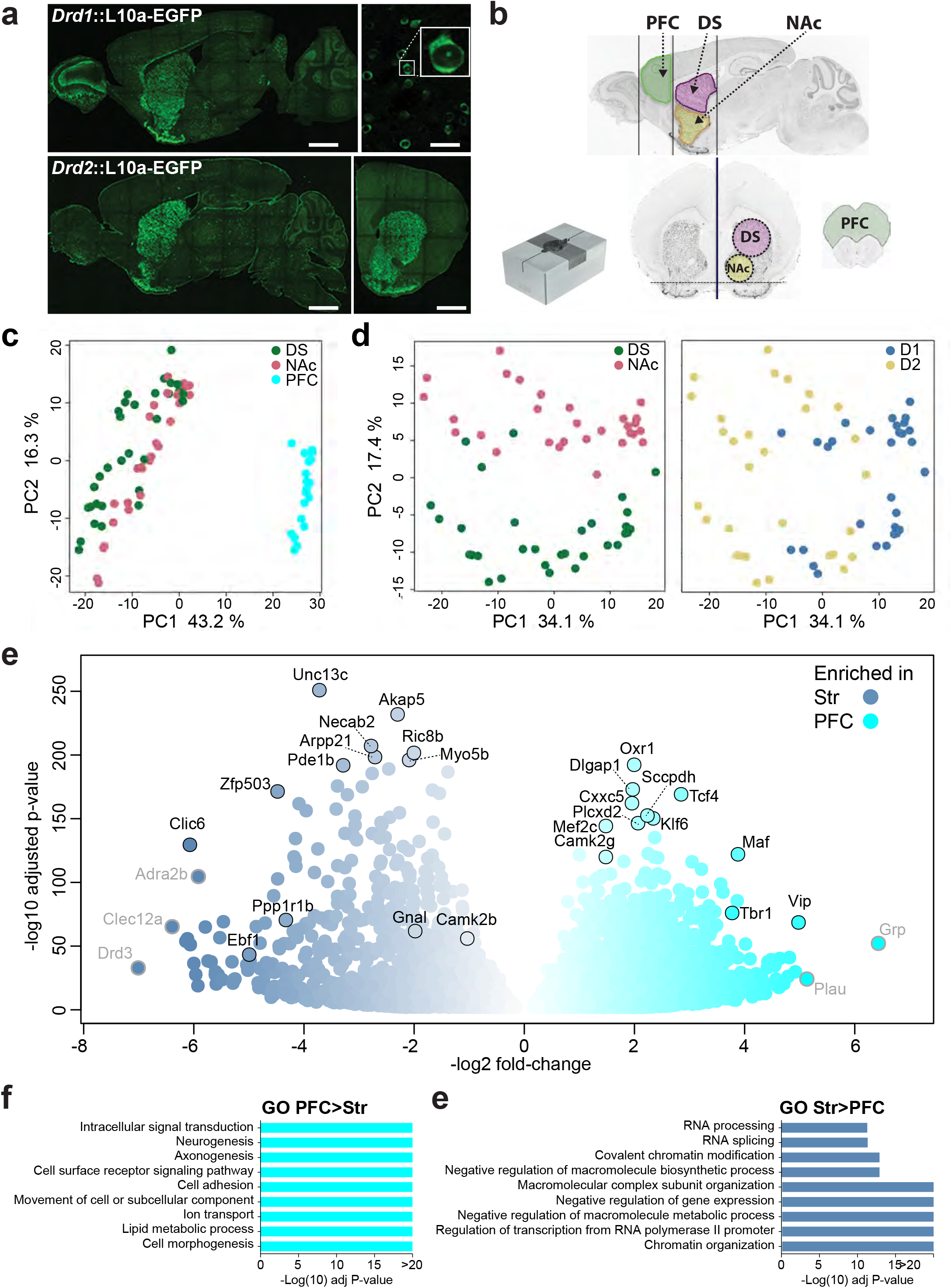
L10-EGFP expression and differences in mRNA expression in the PFC and striatum of TRAP mice. **a** Brain sections from representative TRAP mice showing the location of the cells expressing EGFP-Rpl10a (direct EGFP fluorescence). **Upper panel**, Drd1-EGFP/Rpl10a mouse, left picture sagittal section (scale bar 1.5 mm), right picture higher magnification of the striatum (scale bar 50 μm) and blow up of a single neuron illustrating the cytoplasmic and nucleolar labeling. **Lower panel**, Drd2-EGFP/Rpl10a mouse mice. Left picture, sagittal section, right picture, coronal section through the striatum (scale bars 1.5 mm). Images are stitched confocal sections. **b**. Collection of brain tissue samples. Brains were rapidly dissected and placed in a stainless steel matrix (middle panel with 0.5 mm coronal section interval, and two thick slices containing the **PFC** (green, 2 mm-thick) and the striatum (3 mm-thick) were obtained. The PFC was cut and the dorsal striatum (**DS**, pink) and the nucleus accumbens (**NAc**, yellow) were punched out with a metal cannula on ice (indicated as a dashed circle). Limits of the tissue samples are indicated on sagittal (left panel) and coronal (right) sections. **c**. PCA of RNA-Seq gene expression assessed in TRAP-purified mRNAs from PFC, DS and NAc of Drd1- or Drd2-EGFP/Rpl10a mice. Each point corresponds to a sample of tissues from 1-3 mice. **d**. PCA of RNA-Seq from the DS and NAc of Drd1- and Drd2-EGFP/Rpl10a. The same plot was differentially colored for DS and NAc samples (left panel) or D1 and D2 samples (right panel). **e**. Volcano plots showing the mRNA differential expression between striatal D1 samples (blue) and D1 samples from PFC (cyan). The name of top representative mRNAs are indicated (those with low expression levels are in grey). **f-g**. Main gene ontology (GO) pathways for genes more expressed in PFC than in Str (**f.**) or more expressed in STR than in PFC (**g.**). Only the most significant non-redundant pathways are shown. For complete results see **Supplementary Tables 3g, h**.

RNAseq at high read depth yielded 37-62 million reads per sample (**Supplementary Table 1b**) and a total of 20, 689 genes products from over 25, 883 documented genes were mapped in at least one sample (**Supplementary Table 1c**). Read numbers were low for signature transcripts of non-neuronal cells including astrocytes (although they express DRD1^27^), oligodendrocytes, oligodendrocyte-precursor cells, and microglia (**Supplementary Table 1d**) validating our approach for selecting neurons. Principal component analysis (PCA) of all samples showed that the data were reproducible and that the biological replicates variability was lower than the differences between regions (**Fig. 1c,d**). The main source of variance between the 79 samples (43 %) aligned with the brain region, PFC vs striatum (**Fig. 1c**). Within the striatum, D1 vs D2 cells aligned with PC1 with 34 % of the variance, while DS vs NAc aligned with PC2 with 17 % of the variance (**Fig. 1d**). We then compared the translatomes of these various populations of DA target neurons using DESeq2 (**Supplementary Table 2**). For clarity the results are presented as two-by-two comparisons.

### Comparison of D1 polysome mRNA in PFC and striatum

We first focused on the gene products differentially translated between D1 neurons of the PFC and the striatum (Str, i.e. pooled DS and NAc, **Fig. 1e, Supplementary Tables 3a,b**). Eight thousand gene products were differentially associated with polysomes between these two cell populations, even with a stringent significance threshold (padj < 0.01). This large number of significant differences illustrates the power of TRAP-Seq applied to many independent biological replicates for each region (here 19 D1-PFC, 30 D1-Str, including 15 DS and 15 NAc). It also underlines the existence of major differences between the transcriptional/translational landscapes in cortical and striatal neurons expressing DRD1.

Because many significant differences were of small amplitude and/or corresponded to transcripts with low levels of expression, we applied more filters to identify a limited number of gene products presumably reflecting biologically relevant differences between the two cell populations: padj < 0.001, fold-change ≥ 2 and expression level ≥ 10 counts per million (CPM, **Supplementary Tables 3c,d**). We also selected the mRNAs that were higher in all samples of PFC than in all striatal samples, or vice-versa (**Supplementary Tables 3e,f**). We thus identified a core set of genes with strong and consistent differences between the two regions. These differentially translated genes include, as expected, transcripts considered as characteristic of specific cell populations, such as cortical pyramidal cells or SPNs. Other differences provide information about the distinct properties of cells expressing D1R in PFC and striatum, as illustrated by those with identified functions in the *International union of basic and clinical pharmacology* (IUAPHAR) data base (**Supplementary Tables 3i,j**). Gene ontology (GO) analysis indicated that genes more expressed in PFC are related to neuronal differentiation and morphogenesis, cell adhesion and signaling (**Fig. 1f, Supplementary Table 3g**), whereas those more expressed in Str are related to RNA processing, chromatin, and transcription (**Fig. 1g, Supplementary Table 3h**).

The depth of sequencing and number of samples analyzed allowed us to investigate differences in usage of individual exons, corresponding to different mRNA isoforms generated by alternative splicing and/or selection of transcription start site or polyadenylation site (**Supplementary Table 4**). DEXseq breaks down open reading frames into exon fragments, and tests for significant changes in the relative usage of each of these fragments, i.e. in the ratio of sequencing reads mapping to this fragment, over the reads mapping to any other fragment of the same gene. Approximately 2,000 exon fragments were identified as differentially used (**Supplementary Tables 5a,b**), with several differences often occurring in the same genes (**Supplementary Table 5c**). These exon usage differences were dissociated from those in total gene expression (congruent in only 20-30% of genes with exon usage differences, **Supplementary Tables 5d,e**). A striking example is *Arpp21*, which included 42 exons more used in PFC than in Str and 19 more used in Str than in PFC (**Supplementary Table 5f**). Interestingly, the striatal-enriched exons included the coding sequence of ARPP-21 (**Supplementary Table 5f**, highlighted blue), a regulator of calmodulin signaling^28^ enriched in SPNs^29^, whereas the PFC-enriched exons included those coding for TARPP (highlighted green), a longer protein first described in thymocytes^30^, and now shown to bind RNA through domains absent from the shorter isoform^31^. These results reveal the high degree of cell type specificity of isoform expression, which operates, at least in part, independently of general gene expression regulation. Taken together our data provide the first in-depth the characterization of the transcripts in DRD1-expressing cells in the PFC and striatum and serve as a unique resource for future investigations.

### Comparison of D1 and D2 polysome mRNA in the striatum

We then separately examined the differences between D1 and D2 populations in the DS and in the NAc (**Fig. 2a-c, Supplementary Table 2**). Several thousand differentially expressed genes were identified (padj < 0.01, **Fig. 2c**). Because our TRAP approach could potentially enrich polysomes from both D2-SPNs and cholinergic interneurons (ChIN), which also express *Drd2*^13^, we first examined ChIN markers to evaluate their contribution (**Supplementary Tables 1d** and **2**). The ChIN markers that were detected were significantly enriched in D2 vs D1 neurons but were expressed at very low levels (< 5 CPM, **Supplementary Tables 1d**), indicating that although transcripts from ChINs were immunoprecipitated by *Drd2*-TRAP they represented a minor component of the total mRNA. The low contribution of ChINs in our study contrasts with that observed using D2-Ribo-Tag mice^32^ and may result from the different promoters driving the expression of the reporter in these two methods.

**Fig. 2:**
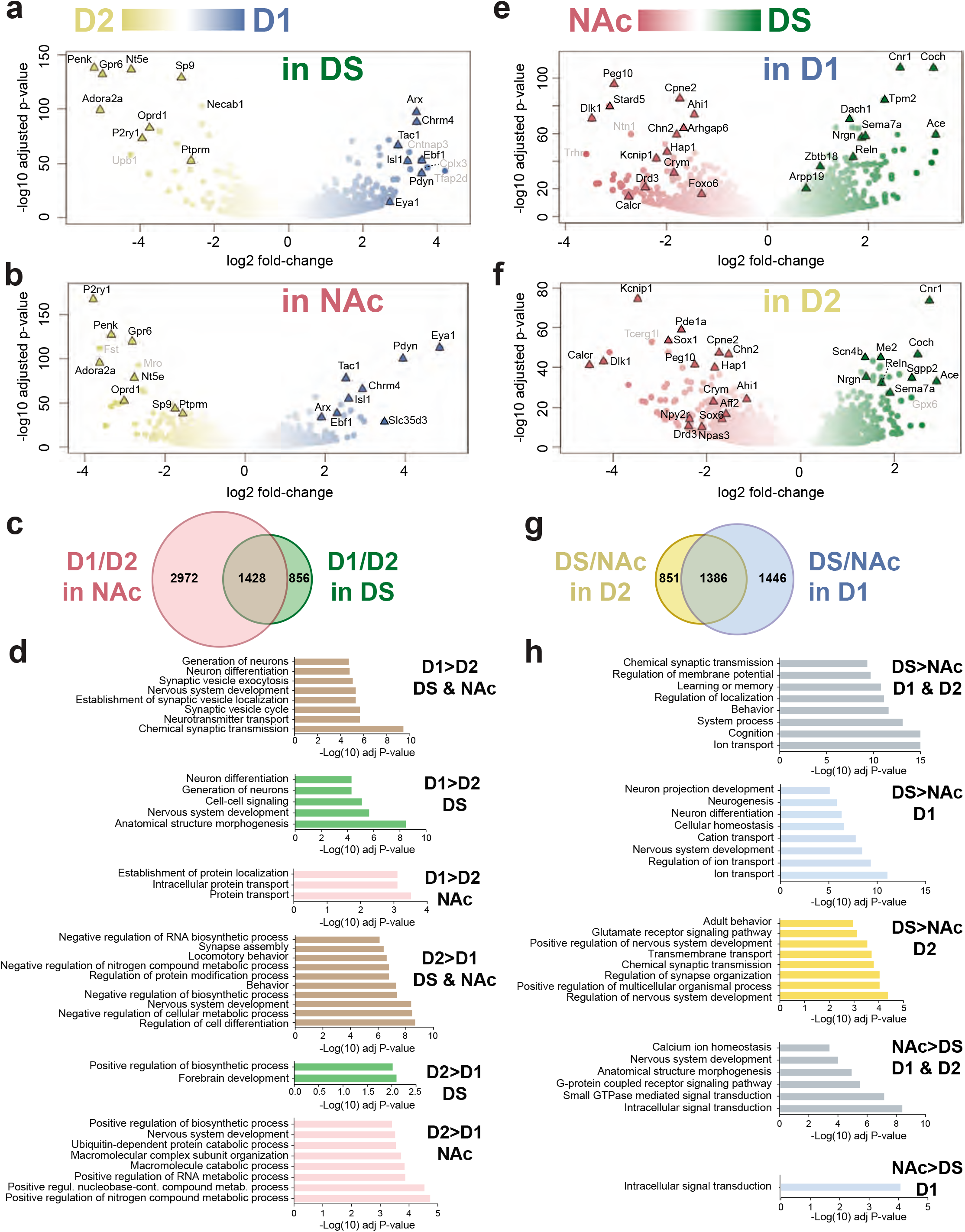
Differential mRNA expression in striatal regions and neuronal populations. mRNA was purified by BAC-TRAP from the DS and NAc of Drd1- or Drd2-EGFP/Rpl10a mice and analyzed by RNA-Seq. **a-b**. Volcano plots of the differences in expression patterns between D1 (blue) and D2 (yellow) samples in the DS (**a**) or in the NAc (**b). c**. Venn diagram of data in a-b showing the number of mRNAs differentially expressed in D1 vs D2 samples in the NAc (light red) and DS (green). **d**. Main gene ontology (GO) pathways for genes more expressed in D1 or in D2 neurons in DS, NAc or both, as indicated. Only the most significant non-redundant pathways are shown. For complete results see **Supplementary Tables 7a-f. e,f**. Volcano plots of the differences between DS (green) and NAc (red) in D1 (**e**) and D2 (**f**) samples. **g**. Venn diagram of the data in **e** and **f** showing the number of mRNAs differentially expressed in DS vs NAc samples in the D1 (blue) and D2 (yellow) samples. In **a, b, d**, and **e** the names of top representative mRNAs are indicated (those with low expression levels are in grey). **h**. Main gene ontology (GO) pathways for genes more expressed in DS or in NAc neurons in D1, D2 or both, as indicated. Only the most significant non-redundant pathways are shown. For complete results see **Supplementary Tables 12a-f**.

We concluded that most of TRAP-seq striatal mRNA originated from SPNs and focused on the major differences between D1- and D2-SPNs, which we analyzed separately in the DS and NAc (**Fig. 2a,b, Supplementary Tables 6a-f**). Importantly many D1/D2 differences were found in both NAc and DS (**Fig. 2c, Supplementary Tables 6a,d**), whereas only 7 genes, with low expression levels, displayed opposite D1/D2 expression differences in DS and NAc (**Supplementary Table 6g**). These results underline the existence of common regulatory mechanisms of gene expression in D1- and D2-SPNs of the DS and NAc.

We also applied the same additional stringent filters as in the PFC/Str comparison above (i.e. fold-change ≥ 2 and expression levels ≥ 10 CPM, **Supplementary Tables 6h-m**, and genes more expressed in all samples of a population than in any sample of the other, **Supplementary Tables 6n-s**) to pinpoint the gene expression differences most likely to have biological significance and provide useful markers. These two complementary strategies identified a set of robust markers largely beyond those previously reported. GO pathways enriched in D1-SPNs as compared to D2-SPNs included chemical synaptic transmission in both DS and NAc, morphogenesis in the DS and protein transport in the NAc (**Fig. 2d, Supplementary Tables 7a-c**). Pathways enriched in D2-SPNs were related to cell differentiation, development, and metabolic processes, and, specifically in the NAc, nucleic acid metabolism and processing (**Fig. 2d, Supplementary Tables 7d-f**). For convenience, differentially expressed genes with identified function (IUPHAR) are shown in **Supplementary Table 7g-h**.

We then examined the D1/D2 differences in exon usage in the DS and NAc (**Supplementary Tables 8,9**). The differences were less numerous in the DS (**Supplementary Tables 10a,b**) than in the NAc (**Supplementary Tables 10c,d**). In either case the same genes often included several differentially used exons (**Supplementary Tables 10e**). Most D1/D2 differences observed in the DS were also found in the NAc, including genes with some exons preferentially expressed in D1 and others in D2 neurons. A characteristic example are the neurexin genes (*Nrx1, Nrx2*, and *Nrx3*), which code for presynaptic adhesion molecules with many splice isoforms and alternative transcription start sites with cell-type specific expression and properties^33^ (**Supplementary Table 10e**, highlighted blue).

### Comparison of DS and NAc polysome mRNA

As shown by PCA (**Fig. 1d**), transcripts markedly differed between DS and NAc in both D1 and D2 neurons, in line with the abundant literature emphasizing the differences between these two regions^16,34^. More than 3,000 genes were differentially expressed between DS and NAC neurons (padj < 0.01, **Figure 2e-f, Supplementary Tables 11a-g**). Strikingly, D1- and D2-SPNs shared many of these dorso-ventral differences (1386, **Figure 2g, Supplementary Tables 11a, d**), whereas very few genes (16) displayed opposite dorso-ventral differences in D1 and D2 neurons (**Supplementary Tables 11g**). As in the PFC/Str and D1/D2 comparisons we applied more filters to identify the genes most likely to be functionally relevant and/or reliable difference markers (**Supplementary Tables 11h-s**, summary of top differentially expressed genes, **Supplementary Table 11t**). GO analysis of differentially expressed genes indicated a predominance of ion transport-related pathways in the DS and signaling in the NAc (**Fig. 2h**, **Supplementary Tables 12d-f**, IUPHAR function in **Supplementary Tables 12g,h**).

We also investigated the DS/NAc differences in exon usage (**Supplementary Tables 13,14**). As above for the D1/D2 differences, these differences were concentrated in a relatively small number of genes which often included several differentially expressed exons (**Supplementary Tables 15a-e**). A notable proportion of DS/NAc differences were common between D1 and D2 neurons (e.g. up to half of those in D1 neurons, **Supplementary Table 15e**) underlining the existence of common regulatory mechanisms in these two populations. Only a small proportion of differences in exon usage corresponded to overall differences in gene expression (**Supplementary Tables 15f-i**).

To expand on the results of exon usage analysis we focused on *Cntnap2*, a gene coding for a cell-adhesion protein, a.k.a. Caspr2, that is associated with autism spectrum disorder and other neuropsychiatric disorders^35^. A short isoform (Iso2) lacks the extracellular domain and corresponding protein-protein interactions of the full-length isoform^36^ (Iso1) (**Fig. 3a, Supplementary Fig. 1a**). Analysis of mRNA expression indicated a relative enrichment of exons specific for Iso2 in the NAc as compared to the DS in both D1 and D2 neurons (**Supplementary Table 15e, Supplementary Fig. 1b,c**).

**Fig. 3:**
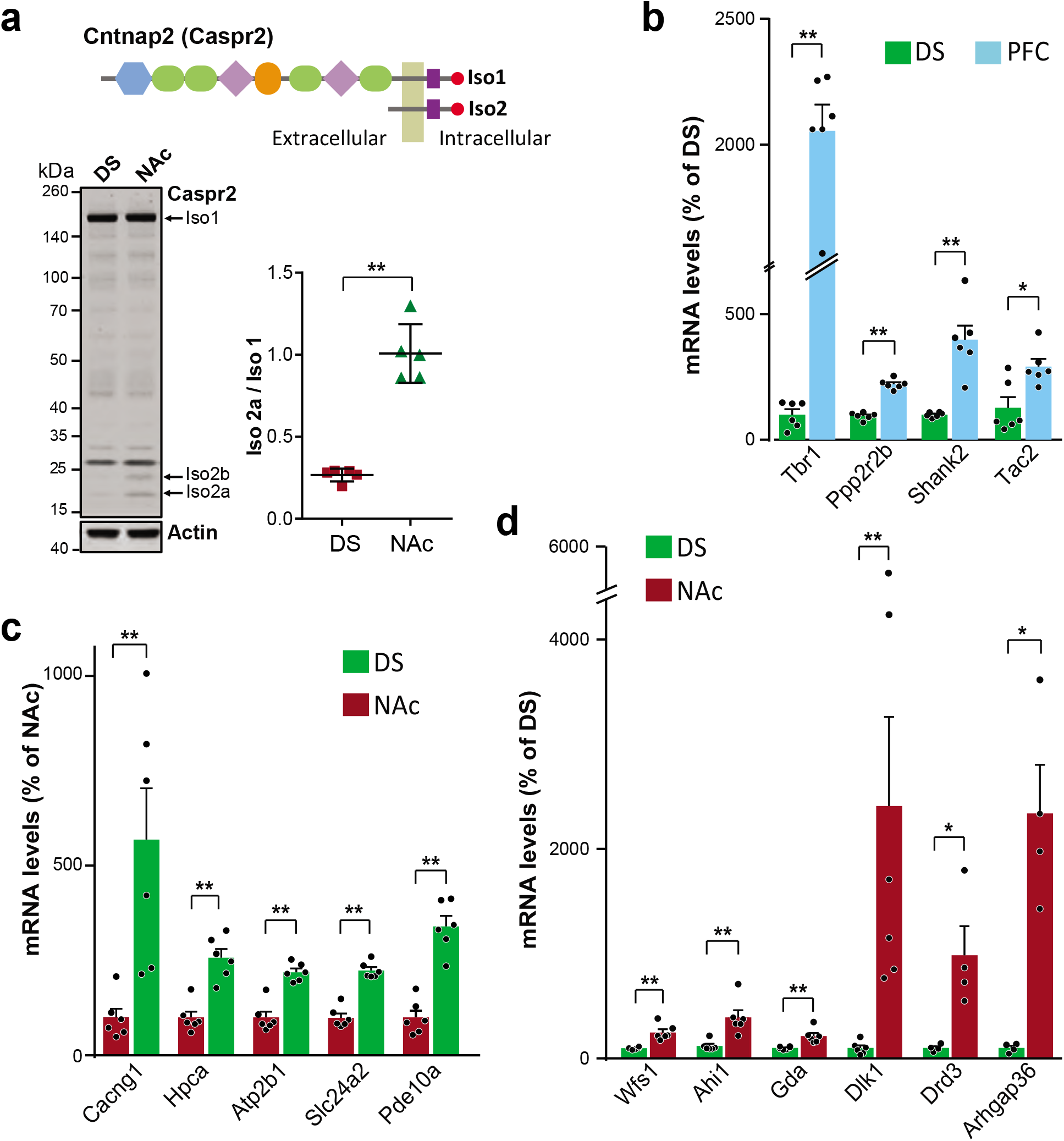
Protein and mRNA expression analyses of selected genes confirming sequencing results. **a**. Immunoblot analysis of the expression of Caspr2 protein coded by *Cntnap2* in mouse DS and NAc. The long (Iso 1) and the short (Iso 2) isoforms are schematized in the top panel (see gene structure in **Supplementary Fig. 2a** and protein domains in Ref^35^). DEXseq mRNA exon usage analysis suggested a higher ratio of Iso2/Iso1 in the NAc than in the DS in D1 and D2 neurons (**Supplementary Fig. 2b,c**). The protein isoform ratio was quantified in samples from 5 mice and normalized to 1 in NAc (means ± SEM are indicated). **b-d**. Genes with diverse levels of differential expression in RNA seq in the regional BAC-TRAP analysis were selected for quantitative PCR verification using wild type mouse brain tissue sample. **b**. Gene products higher in the PFC than in the DS. The expression levels were calculated by the comparative ddCt method and expressed relative to the DS with β-actin as internal control. Data are individual results from independent experiments. Means + SEM are indicated. **c-d**. Same as in (**b**) for selected mRNAs were more expressed DS than in the NAc (**c**) or in the NAc than in the DS (**d**). In **a-d** all statistical analyses were done with two-tailed Mann-Whitney’s test, *p < 0.05; **p < 0.01. Detailed statistical results are presented in **Supplementary Table 19**.

Immunoblotting showed that the Iso2/Iso1 protein ratio was higher in the NAc than in the DS (**Fig. 3a**), confirming at the protein level the TRAP-Seq results. These results suggest possible *Cntnap2* functional differences in NAc and DS in relation to Iso2 levels and illustrate the utility of high resolution translatome comparisons between neuronal populations. Overall the comparison of the NAc and DS separately in D1 and D2 neurons reveals the importance of the dorsoventral differences that, to a large extent, are shared by D1 and D2 SPN populations.

### Validation of results and comparison with other approaches

To assess the validity of the differences observed with TRAP-Seq in independent samples, we carried out reverse transcription followed by real time quantitative PCR (RT-qPCR) on selected transcripts with diverse levels of expression and enrichment using wild type mouse brain tissue samples of mixed cell content. We first confirmed the enrichment of mRNAs in the PFC as compared to the striatum (**Fig. 3b**). We also examined the corresponding *in situ* hybridization patterns available at the Allen Brain Institute (http://mouse.brain-map.org/). For genes with strong enrichment and high expression levels the hybridization differences were striking and consistent with our data (e.g. *Tbr1*) (**Supplementary Fig. 2a**), whereas for others, TRAP-Seq was more informative (e.g. *Ppp2r2b, Shank2, Tac2*). We also confirmed by RT-qPCR the DS/NAc differences for selected genes (**Fig. 3c-d**). Comparison with Allen Brain Institute *in situ* hybridization showed that only some of these differences were visually detectable on available sections (**Supplementary Figs. 2b,c**).

Since the transcriptional patterns in D1 and D2 striatal cell populations have already been explored with various approaches, we compared our results with those previous studies. Although the number of differences we identified was much higher than in a previous work with TRAP followed by microarrays^18^, the amplitudes of differences in gene expression identified in both studies were well-matched (**Supplementary Fig. 3a**). Most of the few genes for which we did not replicate differential expression had low fold-changes in both studies, whereas TRAP-Seq allowed identification of many novel differentially expressed genes (**Supplementary Fig. 3b,c**). We also confirmed 80 % of the D1-associated and 67 % of the D2-associated genes identified in a recent single cell study^21^ and revealed many others (**Supplementary Fig. 3d**). Most of the genes we did not confirm exhibited a low expression (e.g. *Rbp4*) and/or a low fold-change (⍰Log_2_FC⍰ < 1), and often an opposite D1/D2 specificity. Discrepancies may originate from sampling bias or stochastic dropout for the genes with low base counts in single cells. Overall it is important to underline that single cell sequencing and TRAP-Seq provide different types of information that are complementary with consistent results in their area of overlap. TRAP-Seq provides in-depth comparisons and a strong basis for further analysis of identified populations, including with single cell techniques.

### Transcription factor expression and transcriptional networks

In order to identify putative regulators of the SPN populations’ transcriptional profiles, we focused on transcription factor (TF) mRNA differences between regions and cell types. We first examined differential expression of TFs with stringent criteria (**Supplementary Tables 16a-f**). The top differential expression analysis pinpointed some TFs that were previously described in developmental studies. Examples include higher expression in D1-SPNs of DS and NAc of *Isl1* and *Ebf1*, two TFs which govern striatonigral neuron differentiation^37–39^. Conversely *Sp9* was more expressed in all D2-SPNs and *Ikzf2* (*Helios*) in DS D2-SPNs than in D1-SPNs, in agreement with their role in the development of striatopallidal neurons*^40,41^*. These results replicating findings of previous developmental studies validate the capacity of our approach in adult mice to recognize TFs important for development and specification of SPNs. Importantly, we identified many other transcription factors with D1/D2 or DS/NAc differences (**Supplementary Tables 16a-f**), whose role in striatal differentiation has not yet been explored. Some but not all of these TFs have been associated with neuronal development outside of the striatum^42–44^ and our results provide strong incentive for their exploration in the differentiation of SPNs.

To evaluate the potential functional importance of TFs in the regulation of transcriptional profiles in adult striatal neurons, we then used a gene expression-based network inference procedure (see **Methods**). We pruned the network by retaining only the edges that connected either one or two of the TFs differentially expressed between DS and NAc. We focused on TFs differences in D1 cells, only keeping the 1 % edges with the highest scores. Seven TFs passed the filter and six of them formed a connected subnetwork of clusters (**Supplementary Figs. 4,5**). Coloring this subnetwork with relative expression in D1 and D2 populations or in DS and NAc, suggests that different TFs might be involved in cell type or regional specificities. Genes linked to *Nr4a2*, coding for Nurr1, a key TF in the maintenance of DA neurons, and, in the striatum, associated with the development of dyskinesia^45,46^, and *Ebf1*, known for its role in D1-SPN differentiation^37,38^, are strongly differentially expressed between D1 (blue) and D2 (yellow) cells, the first neighbors of *Nr4a2* and *Ebf1* including many D1-specific genes (**Fig. 4a,b, Supplementary Fig. 4**). These results support a role of *Nr4a2* and *Ebf1* in regulating differences between D1 and D2 SPNs in adult mice.

**Fig. 4:**
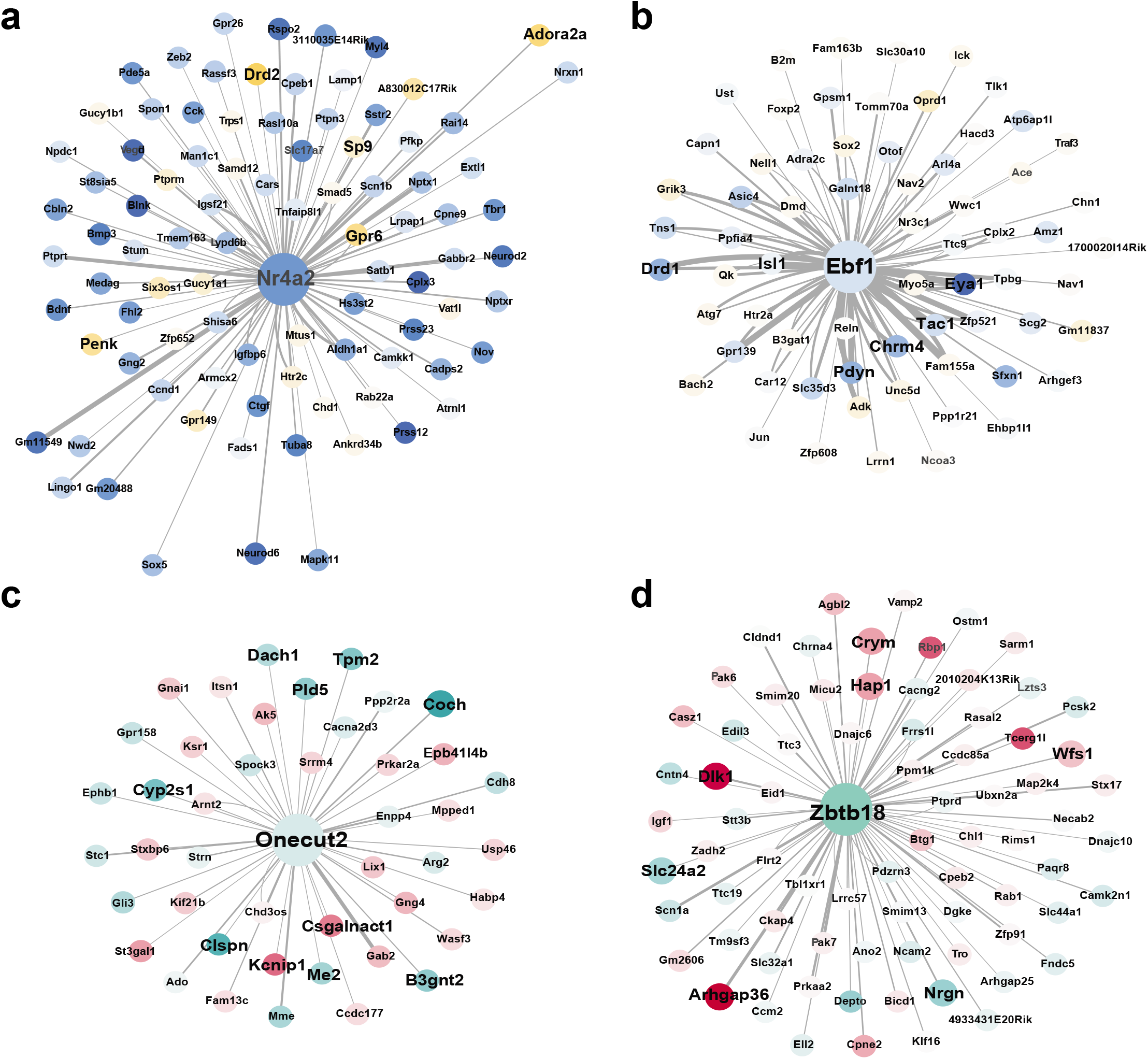
Different transcription factors regulate cellular- and regional-specific gene expression. Subsets of the regulatory network inferred from gene expression pruned using the transcription factors differentially expressed in D1 neurons between DS and NAc, as described in the Methods and Results. Larger networks are shown in **Supplementary Figures 5,6**. Nodes are colored either according to their relative expression in D1 (blue) and D2 (yellow) neurons (**a-b**) or to their relative expression in dorsal (turquoise) and ventral (magenta) striatum (**c-d**). Many genes influencing and influenced by *Nr4a2* (**a**) or *Ebf1* (**b**) are differentially expressed between D1 and D2 neurons. They include markers of D2 and D1 neurons (larger font). In contrast, genes influencing or influenced by *Onecut2* (**c**) or *Zbtb18* (**d**) are more differentially expressed between DS and NAc. Markers of these regions are in larger font (see **Fig. 2** and **Supplementary Tables 6,7**). The color intensities reflect differential expression and the line thickness indicates the reliability of prediction.

In contrast, genes linked to *Onecut2* and *Zbtb18* are strongly differentially expressed in DS (green) and NAc (red, **Fig. 4c,d, Supplementary Figure 5**). In D1 neurons both are enriched in the DS as compared to the NAc (**Supplementary Table 6d**). *Onecut2* is a homeobox gene associated with neuronal differentiation^47^ whose first neighbors include genes enriched in DS (e.g., *Coch, Tpm2, Dach1, Clspn, B3gnt2, Cyp2s1, Pld5, Me2*; **Fig. 4c**). *Zbtb18* mRNA encodes a protein that acts as a transcriptional repressor of key proneurogenic genes and its mutation is implicated in intellectual deficit^48^. *Zbtb18*’s first neighbors include genes enriched in NAc (e.g., *Arhgap36, Wfs1, Crym, Dlk1, Tcerg1l, Hap1*) or in DS (e.g., *Nrgn, Slc24a2*; see **Fig. 4c** and **Supplementary Tables 6a,b,**). Interestingly, among the 83 threshold-passing influences of *Zbtb18*, most (66) are outgoing or bidirectional (7) with a higher weight of outgoing interaction (**Supplementary Fig. 6**), suggesting it is an important upstream regulator of gene expression in the striatum. Thus, our analysis suggests that *Onecut2* and *Zbtb18*, which have not been previously associated with striatal neuron differentiation, could be regulators of dorso-ventral gene expression differences in the striatum.

### Modulatory role of PGE2 in the dorsal striatum

In a different approach to identify candidate upstream regulators that could contribute to the differences between DS and NAc, we used *Ingenuity pathway analysis* (IPA) combining D1 and D2 neurons (**Supplementary Table 17**). Since we noticed that among the endogenous molecules category, prostaglandin 2 (PGE2) was one of the top candidate regulators positively associated with DS-enriched genes, we sought to further investigate the possible role of PGE2 in this region. Little is known about the role of prostaglandins in the striatum, with the exception of a report showing that PGE2 is produced in striatal slices in response to DA receptors stimulation^49^. In that work, the phenotype of mice lacking prostaglandin 1 receptor (PTGER1 a.k.a. EP1, expressed in both D1- and D2-SPNs) indirectly suggested that PGE2 enhanced DRD1 and DRD2 responses^49^. Our mRNA analysis indicated that several genes coding for proteins involved in PGE2 metabolism or action, including PGE2 receptors *Ptger1, Ptger2*, and *Ptger4*, are expressed in SPNs (**Supplementary Table 18**), a result confirmed by single molecule fluorescent *in situ* hybridization (**Fig. 5a-c**). Using RT-qPCR for unequivocal quantitation of transcripts, we found that *Ptger1* and *Ptger2* mRNAs were enriched in the DS as compared to the NAc, especially in D2 neurons (**Fig. 5d,e**).

**Fig. 5:**
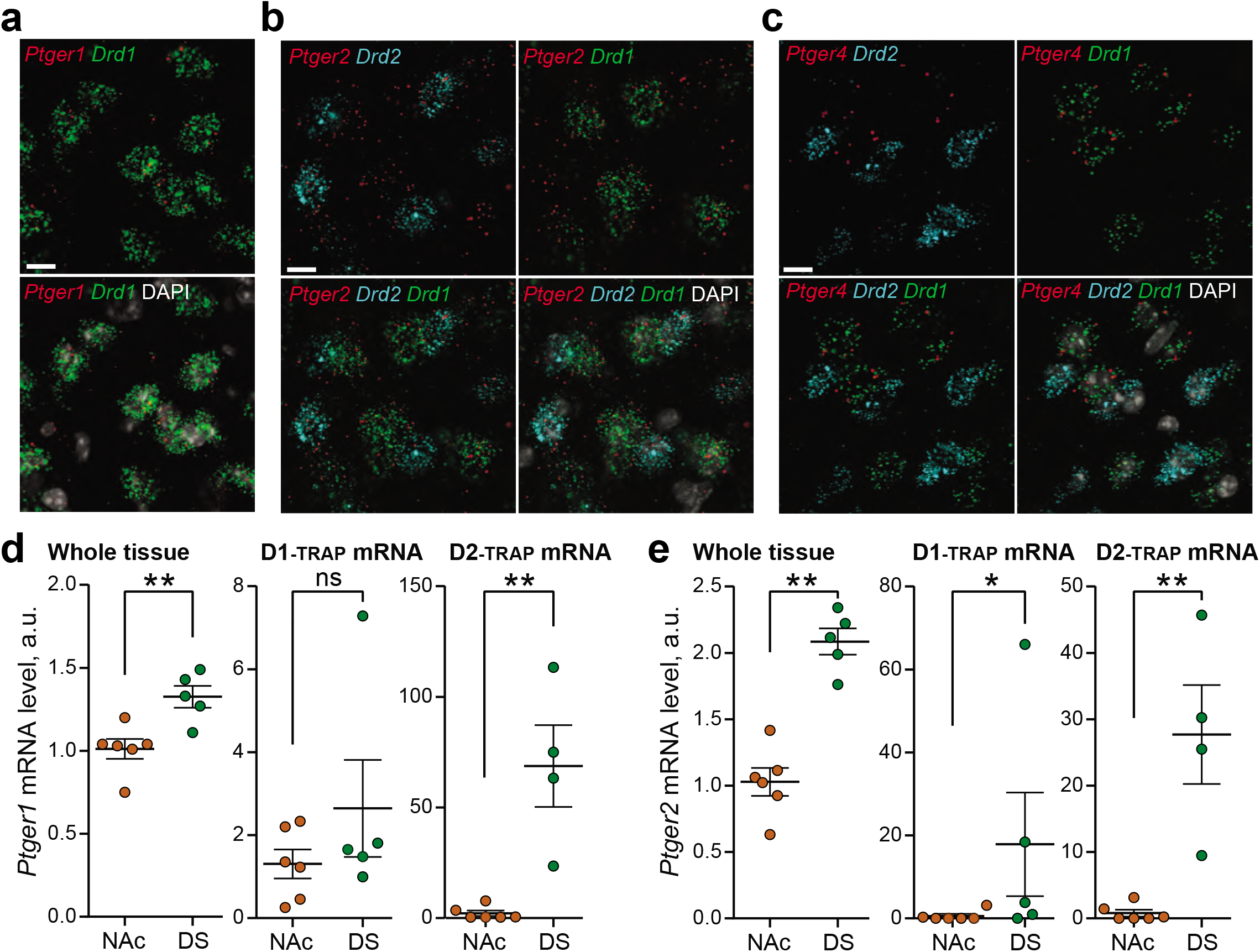
Expression of PGE2 receptors in the striatum. **a.-c. Single molecule fluorescent *in situ* hybridization for PGE2 receptors in the DS**. Sections through the DS of brains from wild-type C57/Bl6 male mice were processed for single molecule fluorescent *in situ* hybridization (smFISH), as described in online Methods. Sections were labeled with probes for one of the PGE2 receptor mRNAs, *Drd1*, and *Drd2*, as indicated, and counterstained with DAPI (grey scale). Images were acquired with a confocal microscope. **a**. smFISH for *Ptger1*. **b**. smFISH for *Ptger2*. **c**. smFISH for *Ptger4*. Scale bars, 10 μm. **d**. and **e**. *Ptger1* (**d**) and *Ptger2* (**e**) mRNA were measured by RT-qPCR in the NAc or DS of wild type mice (left panels), or polysomes purified from *Drd1*- or Drd2-TRAP mice. Expression levels were calculated by the comparative ddCt method and expressed relative to the NAc with rpl19 as internal control. Statistical analysis, Mann-Whitney’s test, statistical significances: * p < 0.05, ** p < 0.01, n.s., not significant (see **Supplementary Table 19** for detailed statistical results). Note that because of gene overlap with *Ptger1* we cannot exclude a contribution of *Pkn1* transcripts.

To pharmacologically stimulate PGE2 receptors in the striatum, we used misoprostol, a non-selective PGE2 agonist, which crosses the blood-brain barrier^50^. To test whether misoprostol i.p. was active in the striatum, we first looked for activation of signaling downstream of Ptger2 (a.k.a. EP2), which is positively coupled to adenylyl cyclase^50^, and measured cAMP-dependent protein phosphorylation by immunoblotting. Thirty minutes after misoprostol injection (0.1 mg.kg^−1^ i.p.), PKA substrate phosphorylation was increased in the DS (**Fig. 6a, Supplementary Fig. 7**), providing evidence for its effects. We then evaluated possible effects on behavior of long-term stimulation of PGE2 receptors, by implanting 12-week-old wild type mice with an osmotic mini-pump delivering misoprostol or vehicle either i.p. or, to exclude peripheral effects, through bilateral cannulas into the DS. We first examined motor performance of these mice in a rotarod test. Mice infused with misoprostol or vehicle, either i.p. or in the DS, learned similarly to remain on an accelerating rotarod (**Supplementary Fig. 8**). However, mice infused with misoprostol in the DS performed better than the vehicle-treated group at a fixed challenging speed (**Fig. 6b**). We cannot exclude that the small amplitude of effects on motor coordination was attributable to the short exposure to PGE2 infusion.

**Fig. 6:**
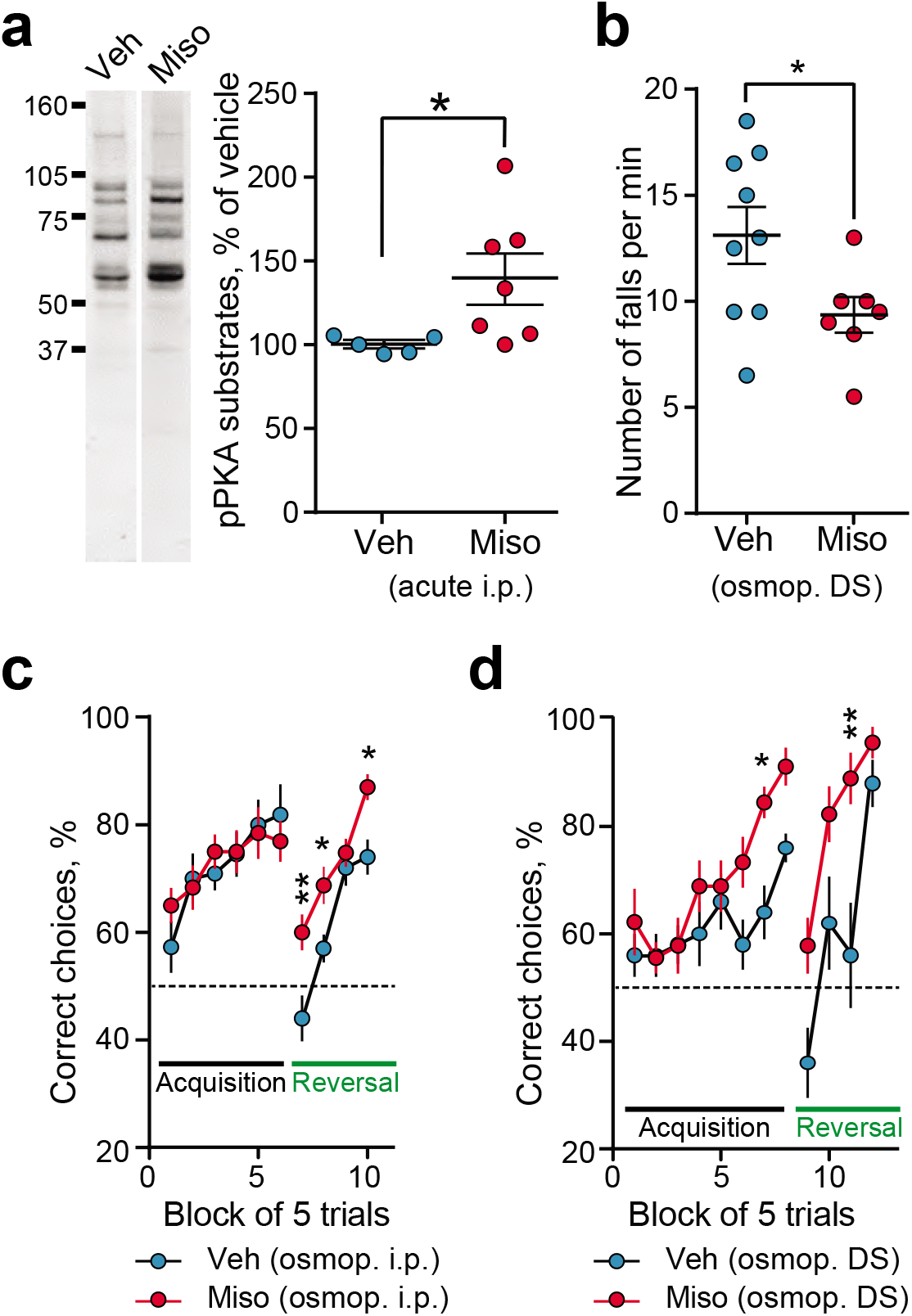
Stimulation of PGE2 receptors effects on signaling, motor behavior and procedural learning. **a**. To test whether the effects of stimulating PGE2 receptors were detected in the DS, misoprostol, a PGE2 agonist, (**Miso**, 0.1 mg.kg^−1^, 7 mice) or vehicle (**Veh**, 5 mice) was injected i.p. Mice were sacrificed 30 min later, striatal extracts were analyzed by immunoblotting with a phosphoPKA-substrate antibody (**left panel** and **Supplementary Fig. 9**). Immunoreactivity was quantified by densitometry and expressed as percent of the mean in vehicle-treated mice (**right panel**). Mann-Whitney’s test (see **Supplementary Table 19). b**. Chronic infusion of a PGE2 agonist in the DS improves rotarod performance. Wild type male with an osmopump delivering Veh (9 mice) or Miso (7 mice) for 9-10 days were trained on accelerating rotarod for 4 days (**Supplementary Fig. 10**) and tested 24 h after the last training session at a fixed speed of 24 r.p.m. Student’s t test (see **Supplementary Table 19). c, d**. Effects of chronic stimulation of PGE2 receptor on procedural learning and reversal. **c**. Wild type male mice were implanted with an i.p. osmopump delivering Veh (20 mice) or Miso (24 mice). Acquisition and reversal of the food-rewarded arm choice in a Y maze was tested 20-25 days later. Two-way ANOVA followed by Holm-Sidak’s post-hoc test (learning phase, n.s., reversal, F_1, 41_ = 17.20, p = 0.0002, *see* **Supplementary Table 19). d**. Same as **c** except that osmopump infusion bilaterally delivered into the DS Veh (10 mice) or Miso (9 mice). Twoway ANOVA followed by Holm-Sidak’s post-hoc test, learning phase, treatment effect, F_(1, 17)_ = 12.17, p = 0.003, reversal, treatment effect, F_(1, 17)_ = 13.62, p = 0.002 (*see* **Supplementary Table 19). a-d**, statistical significances: *p<0.05, **p<0.01, n.s., not significant.

We then evaluated DS-dependent procedural learning in the same mice. We used a food-cued Y-maze, in which mice learn to locate the bated arm with no external cue, using an egocentric strategy^51–53^. The test was carried out in animals receiving misoprostol or vehicle either i.p. or in the DS (**Fig. 6c,d**). The learning phase was similar in mice treated with i.p. infusion of vehicle or misoprostol, but in the reversal task, in which locations of the bated and non-reinforced arms were inverted, relearning was faster in misoprostol-treated mice (**Fig. 6c**). The mice infused in the DS with misoprostol learned faster the stable location of the bated arm and, after reversal, relearned faster than vehicle-infused animals (**Fig. 6d**). Together these results indicate that misoprostol increased procedural learning and its reversal, and that this effect resulted from a local action in the DS.

Because reversal learning is known to be blocked by the injection of DRD2 antagonists in the DS^54^, we hypothesized that PGE2 receptors stimulation might have an opposite effect, mimicking or enhancing DRD2 action. To test this hypothesis and evaluate the functional effects of misoprostol on neuronal activity *in vivo* we used fiber photometry in the DS of awake mice expressing GCaMP6f in D2 neurons (**Fig. 7a-c**). The calcium transients in D2 neurons were increased when mice were exploring a novel environment for 1 min (**Fig. 7a**). This increased activity was observed in mice pretreated with vehicle (**Fig. 7a**), but not in those pretreated with misoprostol (**Fig. 7b**). These results revealed an overall inhibitory effect of misoprostol on D2-SPNs, mimicking stimulation of DRD2 which is known to inhibit these neurons^55^. To test if misoprostol can positively modulate the DRD2 pathway and thus counteract its inhibition, we used a pharmacological approach. We pretreated mice with misoprostol or vehicle before injecting them with haloperidol, a dopamine DRD2 antagonist that induces catalepsy (**Fig. 7d**). Pretreatment with misoprostol inhibited the cataleptic effects of haloperidol, suggesting that stimulation of PGE2 receptors opposes DRD2 antagonist effects. Taken together our results reveal a local modulatory role of PGE2 receptors in the DS, possibly mimicking or enhancing the effects of DRD2 stimulation.

**Fig. 7:**
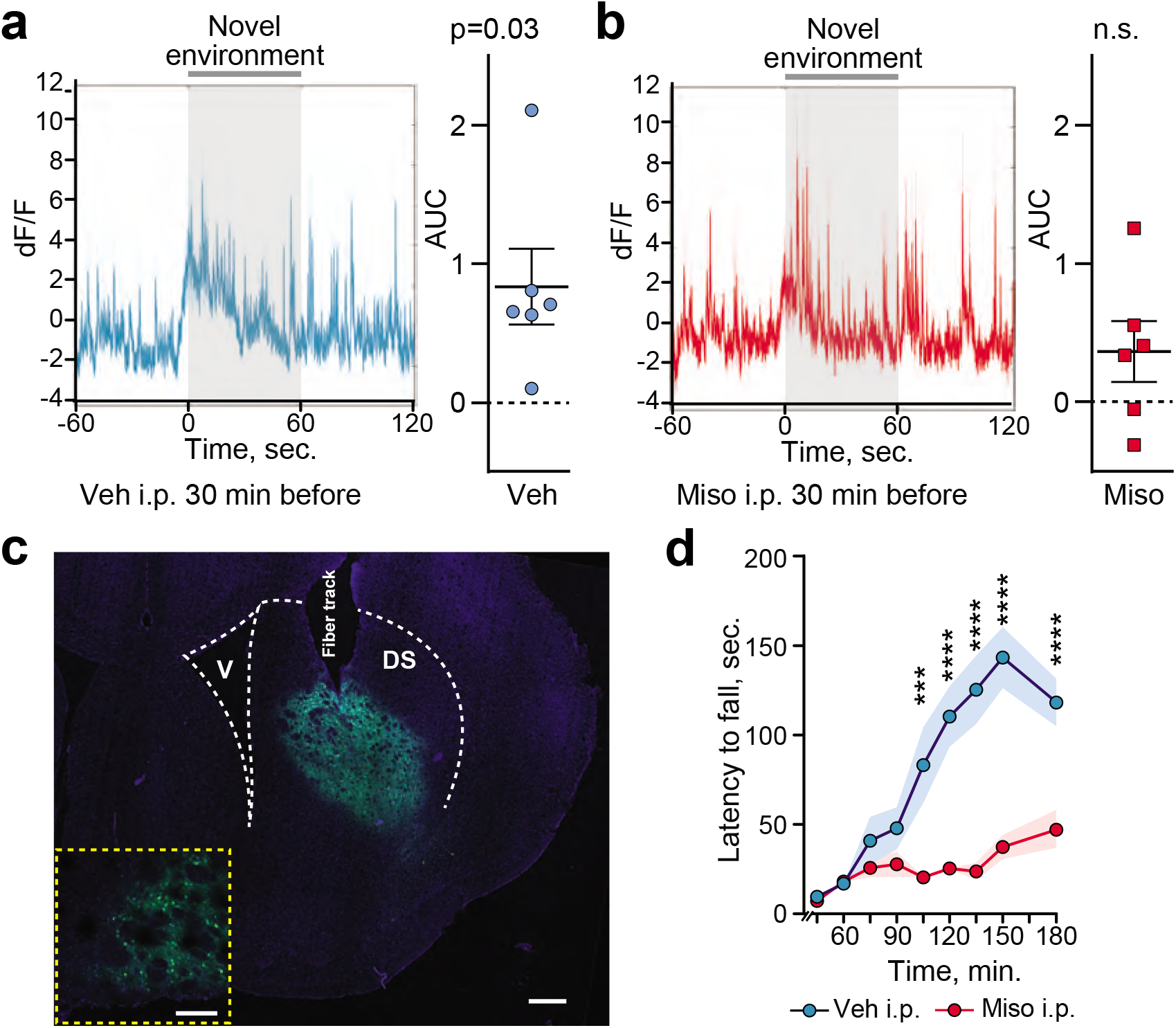
Stimulation of PGE2 receptors decreases D2 neuron Ca^2+^ activity. **a-c**. Acute stimulation of PGE2 receptors decreases D2 neurons activity during exploration of a novel environment. Activity was evaluated by fiber photometry in the dorsal striatum of *Drd2*::Cre mice stereotactically injected with an AAV GCaMP6f (**Supplementary Fig. 11**). Each mouse was recorded twice with an interval ≥ 1 day, 30 min after receiving either vehicle (**Veh**) or misoprostol (**Miso**, 0.1 mg.kg^−1^, i.p.). **a. Left panel:** Representative trace for a mouse injected with vehicle and placed for 1 min in a novel environment, as indicated. **Right panel:** quantification of the area under the curve during novel environment exposure minus baseline for 6 mice. Individual points are shown with mean +/- SEM. Statistical analysis one-sample Wilcoxon test, p = 0.03 (see **Supplementary Table 19). b**. Same as in a. for a mouse injected with misoprostol. One sample Wilcoxon test not significant (n.s.). **c**. Representative histological controls for fiber implantation and GCaMP6f expression. Mosaic of confocal images in a representative mouse used for fiber photometry DAPI (blue)/GCaMP6f (green), scale bar, 200 μm. **Inset:** higher magnification of a different section, scale bar 150 μm. **d**. The effects of PGE2 receptors stimulation on DRD2 function were investigated by evaluating the immobility 45-180 min after haloperidol injection (0.1 mg.kg^−1^, i.p.), in mice pretreated 15 min before with Miso (0.1 mg.kg^−1^, i.p.) or Veh. Two-way ANOVA, post-hoc Holmes-Sidak test (9 mice per group), ***p < 0.001, ****p < 10^−4^, n.s., not significant (see **Supplementary Table 19** for detailed statistical results).

## DISCUSSION

This study reports an in-depth genome-wide regional comparative study of translated mRNAs in the main forebrain dopaminoceptive cell populations expressing DRD1 or DRD2. As expected, striking differences were identified between cortical and striatal DRD1-expressing neurons. In the striatum the differences between the DS and NAc were of a magnitude comparable to those between the D1 and D2 populations. Our work extends previous reports^17–20^ on differences in gene expression between D1- and D2-SPNs and provides the first information about dorso-ventral differences. The use of RNA-Seq instead of microarrays increased > 10-fold the sensitivity of the TRAP approach, based on the number of identified D1/D2 differences. The regional comparisons unveiled previously unexplored differences between the DS and NAc SPNs. Previous analysis of striatal cell diversity by single-cell RNA-seq confirmed the distinction between D1 and D2 SPNs with an additional gradient of transcriptional states attributed to the patch/matrix organization of the striatum^21^. Interestingly, genes defining this gradient correspond to some of those we found highly enriched in the NAc (*Wfs1, Crym*) or the DS (*Cnr1*), suggesting a correlation with the dorso-ventral organization. Although single-cell approaches have the advantage to allow unbiased cell type classification, they are limited to the most highly expressed genes and their interpretation will benefit from comparison with in depth-targeted studies as done here with TRAP-Seq.

Our analyses also provided genome-wide information about exon usage and isoform differences between dopaminoceptive neuronal populations. We found that multiple differences were often detected in the same genes. Most regional or neuronal population exon usage or isoform differences occurred independently from those in global mRNA levels, indicating a dissociation between regulatory mechanisms controlling cell type-specific transcription and those involved in cell type-specific mRNA processing. Our study provides a comprehensive view of the differences between D1- and D2-SPNs, with the first regional comparative evaluation in the DS and NAc. Importantly, we show that for translating mRNA levels as well as for exons usage, many dorso-ventral differences are shared by D1 and D2 neurons, while most D1/D2 differences are found in both the NAc and DS. This reveals the intricacy of regulations of SPN identities, with intersected D1/D2 and DS/NAc influences that give rise to the identity of the various SPN populations. These combined regulations result in regional and SPN-type specificities in information processing at the cellular level, which, together with the differences in anatomical connections, account for their distinct but complementary roles under the modulatory control of DA.

Our analysis of TFs identified potential regulators of these differences between D1/D2 and DS/NAc populations. This approach was validated by the confirmation of the few already known TFs implicated in D1/D2 differences and identified several other novel TFs potentially involved in this function. Among these new TFs candidates for SPN specification, gene network analyses identified a role of *Nr4a2* in D1/D2 differences and of *Zbtb188* and *Onecut2* in DS/NAc differences. These factors, which can now be experimentally investigated during development *in vivo*, may also help refine protocols used to generate specific subtypes of SPNs *in vitro*^56^.

As an example of in-depth regional characterization output, our TRAP results suggested a possible influence of PGE2 in the DS. PGE2 is an important lipid mediator extensively studied outside of the nervous system, but which has previously received little attention in the striatum. We showed the expression of PGE2 receptors in SPNs and explored the potential role of PGE2 using a pharmacological approach. Chronic infusion of misoprostol, a PGE2 agonist, in the DS slightly improved performance on a rotarod test and enhanced procedural learning and its reversal. To address the cellular action of misoprostol we used *in vivo* fiber photometry. We found that an acute injection of the PGE2 agonist inhibited the Ca^2+^ activity of DS D2 neurons during exploration of a novel environment. Moreover misoprostol opposed the cataleptic effects of DRD2 blockade. These results provide converging evidence for the ability of PGE2 to modulate the activity of D2-SPNs *in vivo*, possibly enhancing or mimicking the effects of DRD2 stimulation on these neurons. Our findings on D2 neurons do not exclude PGE2 actions on other cell types which remain to be explored. Because DA is reported to enhance striatal production of PGE2^49^, these effects suggest the existence of a positive PGE2-mediated feedforward regulation DRD2 signaling. Given the key functional role of D2 SPNs^4^, the down-regulation of DRD2 in addiction-like maladaptive behavior^57^ or their sensitivity to neurodegeneration in Huntington’s disease^14^, a potential modulatory role of PGE2 is an exciting possibility that warrants further exploration in future work.

In conclusion, our study provides an extensive in-depth characterization of gene expression and isoform regulation in the main populations of SPNs, comparing them to PFC D1-neurons. It unravels differences between the DS and the NAc and reveals the intersected influences of D1-D2 and dorso-ventral regulations that give rise to SPN subpopulations identities. The comparison of TF expression and network analysis reveals novel transcriptional influences tentatively involved in the striatal regional specification. Our data will help interpretation of results from other approaches and provide means for cell-type specific targeting and therapeutics, as well as a resource for further studies of dopaminoceptive cell functions and genetic risk factors in neuropsychiatric diseases. Moreover, this work reveals an unforeseen role of PGE2 in modulating DS function, especially D2 SPNs, an area with important physiological and pathological implications, open for future exploration.

## METHODS (2974 < 3000)

### Animals

BAC (bacterial artificial chromosome) transgenic mice expressing EGFP fused to the N-terminus of the large subunit ribosomal protein Rpl10a under the control of *Drd1* or *Drd2* promoter (Drd1-EGFP/Rpl10a or Drd1-EGFP/Rpl10a) were previously described^18^, and maintained as heterozygotes on a C57Bl/6J background. Males and females were used (**Supplementary Table 1a**). Wild type male C57Bl/6 mice were purchased from Janvier (France) and used at 10-12 weeks. Mice were maintained on a 12-h light/dark cycle (light off 7:00 pm) and had, before the beginning of the experiment, free access to water and food. Animal protocols were performed in accordance with the National institutes of Health Guide for the Care and Use of Laboratory Animals, and approved by Rockefeller University’s Institutional Animal Care and Use Committee (protocol 14753-H) or in accordance with the guidelines of the French Agriculture and Forestry Ministry for handling animals (decree 87-848) under the approval of the “*Direction Départementale de la Protection des Populations de Paris*” (authorization number C-75-828, license B75-05-22, and APAFIS number 15638, license B751317).

### mRNA extraction

Cell type-specific ribosome-bound mRNA was purified as described^18,25^ with some changes. TRAP transgenic mice were sacrificed by decapitation. The brain was quickly dissected out, placed in cold buffer and then in an ice-cold brain form to cut thick slices from which the PFC was separated and the NAc and the DS punched out using ice-cold stainless steel cannulas (**Fig. 1b**). Samples containing tissue pieces from 1-3 mice (final proportion of males 0.46-0.50 in each cell population, **Supplementary Table 1a**) were homogenized in 1 mL of lysis buffer (20 mM HEPES KOH, pH 7.4, 5 mM MgCl_2_, 150 mM KCl, 0.5 mM dithiothreitol, 100 μg.mL^−1^ cycloheximide (Sigma, #C7698-1g), protease (Roche, #11836170001) and RNAse inhibitors (Ambion, #AM2694, Fisher, #PR-N2515) with successively loose and tight glass-glass 2-mL Dounce homogenizers. Homogenates were centrifuged at 2,000 x g, at 4°C, for 10 min. The supernatant was separated from cell debris, and supplemented with NP-40 (EDM Biosciences, 10 μL.mL^−1^) and 1,2-dihexanoyl-sn-glycero-3-phosphocholine (DHPC, Avanti Polar lipids, 30 mM, final concentrations). After mixing and a 5-min incubation on ice, the lysate was cleared for 10 min at 20,000 x g. A mixture of streptavidin-coated magnetic beads was incubated 35 min at room temperature with biotinylated protein L and then 1 h with EGFP antibody and then added to the supernatant and incubated overnight at 4°C with gentle endover-end rotation. Beads were collected with a magnetic rack, washed 4 times with high-salt buffer (20 mM HEPES-KOH, pH 7.4, 5 mM MgCl_2_, 350 mM KCl, 10 μL.mL^−1^ NP-40) and immediately placed in “RTL plus” buffer (Qiagen). mRNA was purified using RNeasy Plus Micro Kit (Qiagen) and in-column DNAse digestion. RNA integrity was evaluated using a Bioanalyzer with a RNA Pico Chip (Agilent), and the quantity of RNA was measured by fluorimetry using the Quant-IT Ribogreen kit.

### Libraries and sequencing

Five ng of RNA were used for reverse-transcription, performed with the Ovation RNA-Seq System V2 (Nugen). cDNA was quantified by fluorimetry, using the Quant-iT Picogreen reagent, and ultra-sonicated using a Covaris S2 sonicator (duty cycle 10 %, intensity 5, 100 cycles per burst, 5 min). Two hundred ng of sonicated cDNA were used for library construction with the Illumina TruSeq RNA sample prep kit, starting at the End-Repair step, and following the manufacturer’s instructions. The libraries were quantified with the Bioanalyzer high-sensitivity DNA kit, multiplexed and sequenced on an Illumina HiSeq 2500 instrument. At least 20 million 50-bp paired-end reads were collected for each sample.

### Bioinformatics analysis

The quality of the raw data was assessed using FastQC^58^ for common issues including low quality of base calling, presence of adaptors among the sequenced reads or any other overrepresented sequences, and abnormal per base nucleotide percentage. The different libraries were then mapped to the *Mus musculus* genome GRCm38 (UCSC mm10) using HISAT2^59^. After RNA-Seq Quality Control, reads were quantified using the RNA-Seq pipeline of SeqMonk^60^ and, for each gene product, counts of all reads in all exons were exported with the corresponding gene annotations. Gene products from sex chromosomes were not included in the study. Sequencing data has been deposited in NCBI’s Gene Expression Omnibus and are accessible through GEO Series accession number GSE137153 (https://www.ncbi.nlm.nih.gov/geo/query/acc.cgi?acc=GSE137153). Differential expression was performed with DESeq2 package^61^ v1.18 with the option betaPrior set to TRUE. Significance was set at an adjusted p-value of 0.01. Except for differential expression analysis, the raw count files were processed to generate counts per million reads (CPM). Data were then normalized with DESeq2’s rlog function. PCA was performed with the prcomp function of the stats R package. GO term enrichment (biological process branch) was performed on lists of significantly differentially expressed genes with WebGestalt, using the list of all expressed genes as background. Only categories with a FDR of 0.01 after Benjamini-Hochberg correction^62^ were retained.

To analyze differential exon usage across cell populations, we performed more sample filtering. The technology used for reverse transcription and amplification of TRAP RNA (Nugen Ovation RNAseq v2 kit) causes a slight 3’ bias across the body of every gene, which could mask some of the exon usage differences. We computed average read coverages across gene bodies for every sample using the RSeQC package^63^ and then calculated the ratio of read coverage across the 3’-most 20 % of the gene body over coverage across the 5’-most 20 % of the gene body. The largest ratios flag the samples with the largest 3’-bias. We thus filtered out the 10 % of the samples with the largest 3’ to 5’ ratios. For the remainder of the samples, we used the DEXseq package^64^ and the Ensembl release 70 annotations to calculate read counts for each exon in each sample. We loaded the count matrix into R and estimated the size factors, dispersions and Exon Fold Changes by using the DEXSeq package with the default parameters.

### Network inference

Following the conclusions of the DREAM5 challenge’s analysis^65^, we used a combination of methods based on different algorithms to infer regulatory networks. We combined the results of CLR^66^, a mutual-information-based approach providing undirected edges, and GENIE3^67^, a tree-based regression approach providing directed edges. Both methods were the best performers in their category at DREAM5. Both tools were applied using gene expression from all samples, using filtered CPM as described above. Because CLR only provides undirected edges while GENIE3 provides directed ones, the results of CLR were all mirrored with the same score on edges in both directions. Only edges with a positive score in GENIE3’s results were used. The edges present in both CLR and GENIE3 results were then ranked according to the product of CLR and GENIE3’s scores. This only retains edges that have either an extremely high score with one method, or consistent scores with both methods. Subnetworks were extracted using gene lists as seeds, retaining only the first neighbors with scores above a threshold. Visualization and analysis of the resulting networks was done using Cytoscape^68^. The R script is available as <u>Supplementary material</u>: *Network-Inference.R*

### Total RNA purification, cDNA preparation and real-time PCR

Tissue samples (1 mouse per sample, dissected as above) were homogenized in TRIzol with loose and tight glass-glass 2-mL Dounce homogenizers. Total RNA was extracted with TRIzol Reagent (Life Technologies) according to manufacturer’s instructions. The RNA was quantified with a Nanodrop 1000 spectrophotometer and its integrity checked with the Bionalyzer (Agilent RNA 6000 nano kit). RT-qPCR, was performed using SYBR Green PCR kit in 96-well plates according to the manufacturer’s instructions. For analysis of cell-specific mRNA, 5 ng of RNA were used for reverse-transcription, performed with the Ovation RNA-Seq System V2 (Nugen). qPCR was performed in a LightCycler 1.5 detection system (Roche, Meylan France) using the LightCycler FastStart DNA Master plus SYBR Green I kit (Roche). Results were quantified and normalized to a house-keeping gene with delta-delta-CT (ddCT) method.

### Single molecule fluorescent *in situ* hybridization

Analyses of *Ptger1, Ptger 2, Ptger 4, Drd2*, and *Drd1* mRNA expression were performed using single molecule fluorescent *in situ* hybridization (smFISH) as described^32^. Brains from 2 C57/Bl6 male mice were rapidly extracted, snap-frozen on dry ice and stored at −80°C until use. Fourteen-μm coronal sections of the DS (bregma +0.86 mm) were collected directly onto Superfrost Plus slides (Fisherbrand). RNAscope Fluorescent Multiplex labeling kit (ACDBio, #320850) was used to perform the smFISH assay according to manufacturer’s recommendations. Probes used for staining were Mm-Ptger1-C3 (ACDBio, #551308-C3), Mm-Ptger2 (#546481), Mm-Ptger4 (#441461), Mm-Drd1 (#461908), Mm-Drd2-C3 (#406501-C3) and Mm-Drd1-C2 (#461901-C2). After incubation with fluorescent-labeled probes, sections were counterstained with DAPI and mounted with ProLong Diamond Antifade medium (Thermo-Fisher, #P36961). Confocal microscopy and image analyses were carried out at the Montpellier RIO imaging facility. Double- and triple-labeled images were single confocal sections captured using sequential laser scanning confocal microscopy (Leica SP8).

### Pharmacological treatments

For acute treatments, misoprostol (Santa Cruz Biotechnology, Santa Cruz, CA, #SC-201264) was dissolved in phosphate-buffered saline (PBS) and injected i.p. (0.1 mg.kg^−1^). PBS was used as vehicle treatment in control mice. Haloperidol (Tocris) was dissolved in saline and injected i.p. (0.5 mg.kg^−1^). For intra-peritoneal misoprostol infusion, 12-week-old WT mice (n = 23-24 per treatment) were deeply anesthetized with pentobarbital (40 mg.kg^−1^) and an osmotic mini-pump was intraperitoneally (i.p.) implanted (model 1004; Alzet, Palo Alto, CA). Vehicle (DMSO in PBS, 1/1) or misoprostol (Santa Cruz Biotechnology, #SC-201264), were infused at a dose of 50 μg.kg^−1^.day^−1^. For intra-striatal misoprostol infusion, 12-week-old WT mice (n = 9-10 per condition) were deeply anesthetized with pentobarbital (40-60 mg.kg^−1^) and placed in a stereotaxic apparatus for bilateral insertion of a 28-gauge stainless steel cannula (Plastics One, #3280PD-4.0-SPC) into the dorsal striatum (+0.6 mm anteroposterior to bregma, ±2.0 mm lateral to midline and −3 mm ventral to the bone surface). Cannulas were fixed on the skull with anchor screws and dental cement. Osmotic minipumps (model 2004; Alzet, Palo Alto, CA, USA), previously equilibrated overnight at 37°C in PBS, were implanted subcutaneously in the back of the animal and connected to the cannulas allowing infusion of vehicle (PBS) or misoprostol (0.11 μl.h^−1^), resulting in a misoprostol dose of 0.03 μg.day^−1^ per side.

### Immunoblotting

To analyze CNTNAP2/Caspr2 isoforms, DS and NAc were dissected from adult mouse brains and immediately frozen on dry ice. Tissue samples were lysed in RIPA buffer (50 mM Tris, pH 8.0, 150 mM NaCl, 5 mg.mL^−1^ sodium deoxycholate, 10 μL.mL^−1^ NP-40, 1 mg.mL^−1^ SDS) with Complete protease inhibitors (Roche). Equal amounts of protein (BCA protein assay, Thermo-Fisher, #23235) were separated by SDS-PAGE on NuPAGE 8-12 % Bis-Tris gels (Thermo-Fisher) and transferred to nitrocellulose membrane (0.45 mm) in 25 mM Tris-HCl, pH 7.4, 192 mM glycine and 200 mL.L^−1^ ethanol. Membranes were blocked with 50 g.L^−1^ non-fat dry milk in Tris-buffered saline (TBS; 0.25 M Tris, 0.5 M NaCl, pH 7.5) with Tween-20 1 mL.L^−1^ (TBST) for 1 h at room temperature, incubated with previously described^69^ rabbit antibody 187 directed against the intracellular domain of Caspr2, in the same buffer for 2 h and then 1 h with appropriate IRDye-conjugated secondary antibodies. Blots were quantified with the Odyssey–LI-COR infrared fluorescence detection system (LI-COR). For analysis of striatal phosphoPKA-substrate, mice were sacrificed by cervical dislocation 30 min after i.p. injection of misoprostol (0.1 mg.kg^−1^) or vehicle, and striata dissected as above. Fifteen μg of protein per sample were separated in 4-20% SDS-polyacrylamide gel (BIO-RAD mini-protean-TGX) before electrophoretic transfer onto a nitrocellulose blotting membrane (Amersham, #G9990998). Membranes were blocked 45 min in 30 g.L^−1^ bovine serum albumin and 10 g.L^−1^ non-fat dry milk in 0.1 M PBS. Membranes were then incubated overnight with anti-p(Ser/Thr) PKA substrates (1:1000; Cell Signaling Technology, Beverly, MA, USA) and then IRDye-conjugated secondary antibodies, visualized with Odyssey–LI-COR. The optical density in the lane after acquisition was assessed using the GELpro32 software. Results were normalized to the detection of pThr197PKAc (1:1000; Cell Signaling Technology, Beverly, MA, USA) in the same sample.

### Fiber photometry

Male Drd2-Cre mice were anaesthetized with isoflurane and received 10 mg.kg^−1^ intraperitoneal injection (i.p.) of Buprécare^®^ (buprenorphine 0.3 mg) diluted 1/100 in NaCl 9 g.L^−1^ and 10 mg.kg^−1^ of Ketofen^®^ (ketoprofen 100 mg) diluted 1/100 in NaCl 9 g.L^−1^, and placed on a stereotactic frame (Model 940, David Kopf Instruments, California). pAAV.Syn.Flex.GCaMP6f.WPRE.SV40 virus (500 nL, titer ≥ 1X1013 vg/mL, working dilution 1:5, Addgene, #100833-AAV9; https://www.addgene.org/100833/RRID:Addgene_100833) was injected unilaterally into the DS (L = ±1.5; AP = +0.86; V = −3.25, in mm) at a rate of 0.05 μl.min^−1^. The injection needle was carefully removed after 5 min waiting at the injection site and 2 min waiting half way to the top.

A chronically implantable cannula (Doric Lenses, Québec, Canada) composed of a bare optical fiber (400 μm core, 0.48 N.A.) and a fiber ferrule was implanted 100 μm above the location of the viral injection site (L = ±1.5; AP = +0.86; V = −3.25, in mm). The fiber was fixed onto the skull using dental cement (Super-Bond C&B, Sun Medical). Real time fluorescence emitted from GCaMP6f-expressing neurons was recorded using fiber photometry as described^70^. Fluorescence was collected using a single optical fiber for both delivery of excitation light streams and collection of emitted fluorescence. The fiber photometry setup used 2 light-emitting LEDs: 405 nm LED sinusoidally modulated at 330 Hz and a 465 nm LED sinusoidally modulated at 533 Hz (Doric Lenses) merged in a FMC4 MiniCube (Doric Lenses) that combines the 2 wavelengths excitation light streams and separate them from the emission light. The MiniCube was connected to a Fiberoptic rotary joint (Doric Lenses) connected to the cannula. A RZ5P lock-in digital processor controlled by the Synapse software (Tucker-Davis Technologies, TDT, USA), commanded the voltage signal sent to the emitting LEDs via the LED driver (Doric Lenses). The light power before entering the implanted cannula was measured with a power meter (PM100USB, Thorlabs) before the beginning of each recording session. The irradiance was ^~^9 mW.cm^−2^. The fluorescence emitted by GCaMP6f in response to light excitation was collected by a femtowatt photoreceiver module (Doric Lenses) through the same fiber patch cord. The signal was received by the RZ5P processor (TDT). Real time fluorescence due to 405-nm and 465-nm excitations was demodulated online by the Synapse software (TDT). A camera was synchronized with the recording using the Synapse software. Signals were exported to MATLAB R2016b (Mathworks) and analyzed offline. After careful visual examination of all trials, they were clean of artifacts in these time intervals. The timing of events was extracted from the video. For each session, signal analysis was performed at two time intervals: −60 to 0 sec (home cage) and 0 to +60 s (new environment, NE). From a reference window (from −180 to −60 sec), a linear leastsquares fit was applied to the 405 nm signal to align it to the 465 nm signal, producing a fitted 405 nm signal. This was then used to calculate the ΔF/F that was used to normalize the 465 nm signal during the test window as follows: ΔF/F = (465 nm signaltest - fitted 405 nm signalref)/fitted 405 nm signalref. One sample Wilcoxon test was performed on values from the AUC calculated during NE (0 to +60 s) minus the AUC during the home cage (−60 to 0 s) for each trial. Each mouse was recorded twice with an interval of at least a day and received an i.p. injection of misoprostol (0.1 mg.kg^−1^) or vehicle 30 min before the start of the recording. The first time, the treatment was chosen randomly and the second time, mice received the other treatment.

### Behavioral assays

#### Haloperidol-induced catalepsy

mice were injected with misoprostol (0.1 mg.kg^−1^, i.p.) or vehicle, and 15 min later with haloperidol (0.5 mg.kg^−1^, i.p.). Catalepsy was measured at several time points, 45-180 min after haloperidol injection. Animals were taken out of their home cage and placed in front of a 4-cm-elevated steel bar, with the forelegs upon the bar and hind legs remaining on the ground surface. The time during which animals remained still was measured. A behavioral threshold of 180 sec was set so the animals remaining in the cataleptic position for this duration were put back in their cage until the next time point.

Behavior of mice chronically implanted with osmotic minipumps was explored successively using rotarod and food-cued Y maze, 9-15 days and 20-25 days after implantation, respectively.

#### Rotarod

Animals were placed on a motorized rod apparatus (30-mm diameter) accelerating linearly from 4 to 40 RPM over 5 min. Training was performed 14 times (4 trials per day) over consecutive days. The fall latency was recorded. Twenty-four h after the last training session the animals were tested at fixed speed (24 RPM) and the number of falls per min was counted.

#### Y-maze

Mice were tested for learning and cognitive flexibility in a black Y maze (arm 35-cm length, 25-cm height, 15-cm width). All mice were mildly food deprived (85-90 % of original weight) for 3 days prior to starting the experiment. The first day mice were placed in the maze for 15 min for habituation. Then, mice underwent 3 days of training with one arm reinforced with a highly palatable food pellet (TestDiet 5-UTL). Each mouse was placed at a start point and allowed to explore the maze. It was then blocked for 20 sec in the explored arm and then placed again in the starting arm. This process was repeated 10 times per day. At the end of the learning phase all mice showed a > 70 % preference for the reinforced arm. The average number of entries in each arm over 5 trials was plotted. Two days of reversal learning followed the training phase during which the reinforced arm was changed and the mice were subjected to 10 trials per day with the reward in the arm opposite to the previously baited one.

## Supporting information

List of supplementary Tables

## ENDNOTES

## Acknowledgements

The manuscript is dedicated to the memory of Paul Greengard who passed away on April 13^th^, 2019, before the manuscript was completed.

Authors thank the Babraham Institute’s Bioinformatics team for help with read mapping and counting, Vincent Knight-Shrijver for his volcano-plot R script, Lucile Marion-Poll for helpful suggestions, and Serge Luquet for his support for the experiments carried out at the BFA.

## Authors contributions

JAG and JPR conceived and supervised the project. EM, JPR, AG, CM, LG, EV, DH, NGlN, and JAG designed the experiments. EM, JPR, AG, YN, CM, BdP, AP, LG, LC, and EV performed experiments. EM, JPR, AG, CM, LG, ACN, EV, DH, and JAG analyzed data, LT, WW, KDN, LV, NGlN, and JPR analyzed and interpreted bioinformatics analyses, EM, AG, LT, YM, CM, BdP, AP, LG, ACN, EV, DH, NH, NGlN, PG, and JAG discussed the data and provided input and corrections to the manuscript. EM, JPR, and JAG wrote the manuscript. All the authors but PG approved the final version of the manuscript.

## Conflict of interest

The authors declare no conflict of interest.

## Support

This work was supported by Inserm and Sorbonne Université, and grants from European Research Council (ERC, AIG-250349) and Biology for Psychiatry Laboratory of excellence (Labex Bio-Psy) to J.A.G., *Fondation pour la Recherche Médicale* (FRM # DPA20140629798) and ANR *Epitraces* (Project-ANR-16-CE16-0018) to J.A.G. and E.V., ANR-17-CE37-0007 (Metacognition) to C.M., the United States Army Medical Research and Material Command (USAMRMC) Award No. W81XWH-14-1-0046 to J.P.R., the Fisher Center for Alzheimer’s Disease Research to P.G., NIH grants DA018343 and DA040454 to A.C.N. E.M. was supported by a Marie Curie International Training Network (ITN) N-PLAST. A. G. is a Ramón y Cajal fellow (RYC-2016-19466) and is supported by a grant from the *Ministerio de Ciencia, Innovación y Universidades* (Spain) (Project no. RTI2018-094678-A-I00). Y.N. was recipient of a Uehara Memorial Foundation fellowship and a Fyssen Foundation fellowship. NGL was supported by BBSRC (BB/P013406/1, BB/P013414/1, BB/P013384/1). KDN received an Amgen Scholarship. LV received support from the Erasmus+ program.

## Data sets

Sequencing data have been deposited in NCBI’s Gene Expression Omnibus and are accessible through GEO Series accession number GSE137153 https://www.ncbi.nlm.nih.gov/geo/query/acc.cgi?acc=GSE137153

## Notes

### Competing Interest Statement

The authors have declared no competing interest.

