## Supplementary Figures for "Translational profiling of mouse dopaminoceptive neurons reveals a role of PGE2 in dorsal striatum"

Enrica Montalban et al.

### SUPPLEMENTARY FIGURES

### List of Figures:

- **Supplementary Figure 1:** Comparison of exon usage for Cntnap2 mRNA in DS and NAc of D1 and D2 neurons.
- **Supplementary Figure 2:** Examples of *in situ* hybridization for differentially expressed mRNAs in PFC DS and NAc.
- **Supplementary Figure 3:** Comparison of the results of the current study with previous reports on D1 D2 differences.
- **Supplementary Figure 4:** Striatal mRNA network of transcription factors illustrating connections with mRNAs differentially expressed in D1 and D2 cells.
- **Supplementary Figure 5:** Striatal mRNA network of transcription factors illustrating connections with mRNAs differentially expressed in DS and NAc.
- **Supplementary Figure 6:** mRNA expression-based network inference of Zbtb18 regulations in SPNs.
- **Supplementary Figure 7:** Immunoblots of pPKA-substrates quantified in Fig. 6a.
- **Supplementary Figure 8:** Lack of effect of a prostaglandin agonist on rotarod learning.

- **References for Supplementary Figures**

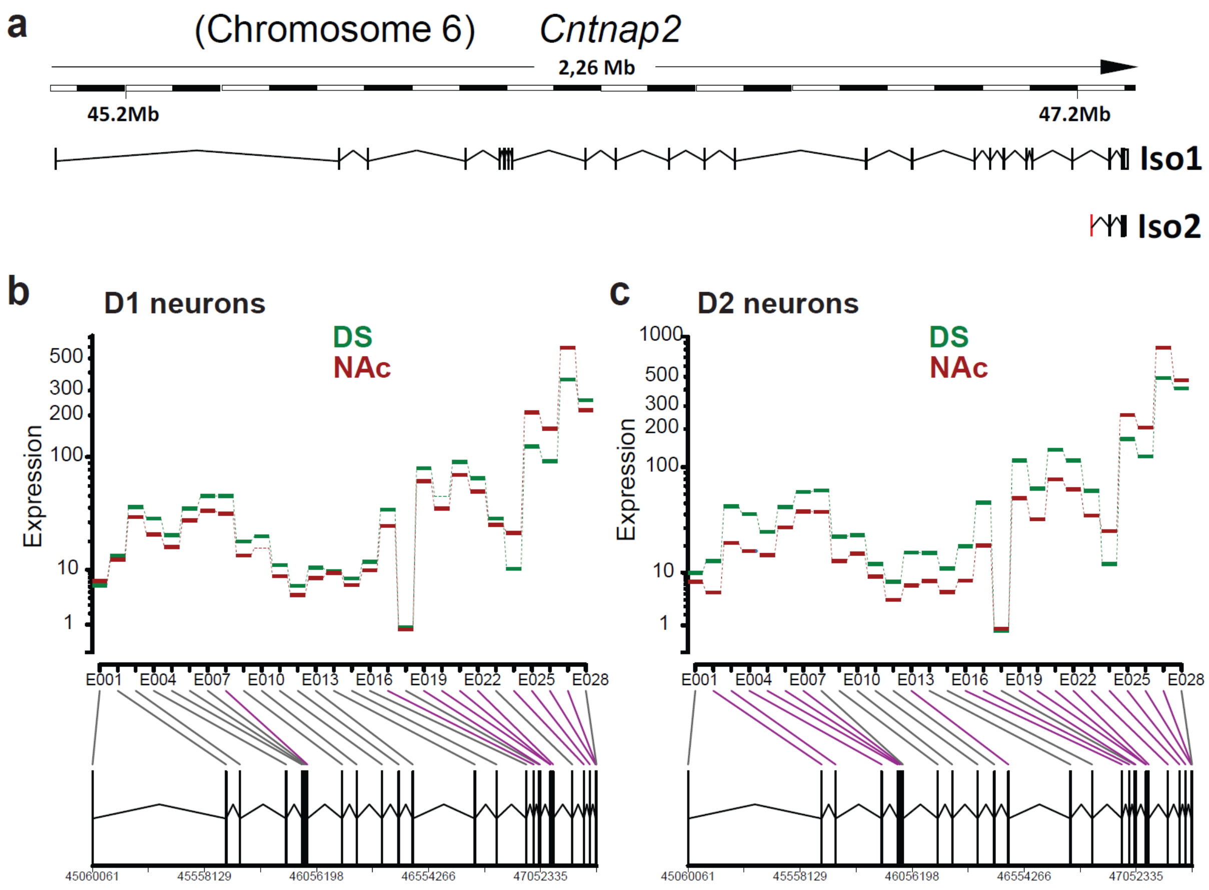


**Supplementary Figure 1: Comparison of exon usage for *Cntnap2* mRNA in DS and NAc of D1 and D2 neurons.**

**a.** Structure of the *Cntnap2* gene which codes for the protein Caspr2. Iso2 is generated by the use of an internal promoter and exon 24. **b.** DEX-Seq plot of the relative expression levels of the various exons of *Cntnap2* in DS and NAc of Drd1-EGFP/Rpl10a mice. **c.** DEX-Seq plot of the relative expression levels of the various exons of *Cntnap2* in DS and NAc of Drd2-EGFP/Rpl10a mice.

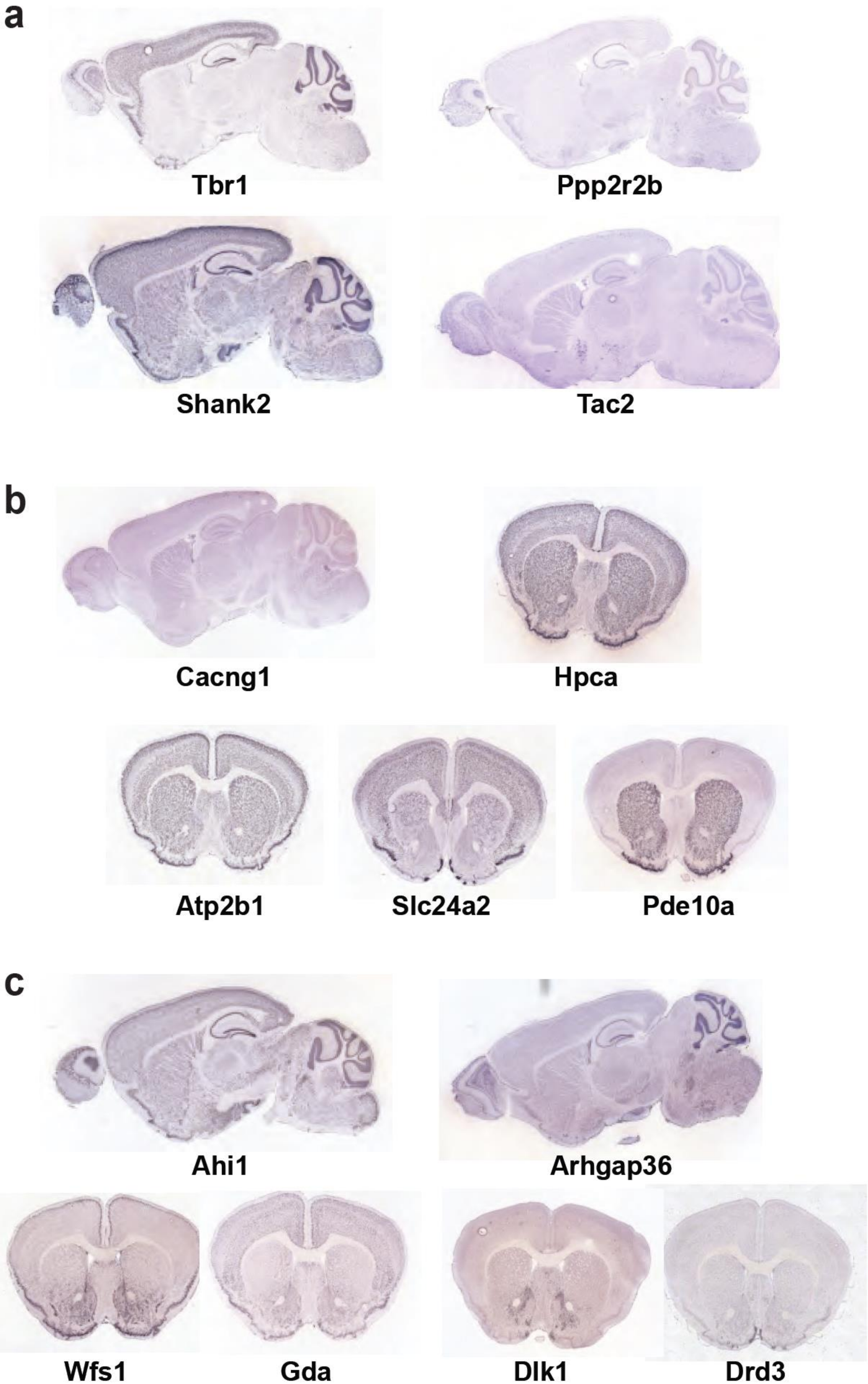
a
Tbr1
Ppp2r2b
Shank2
Tac2
b
Cacng1
Hpca
Atp2b1
Slc24a2
Pde10a
c
Ahi1
Arhgap36
Wfs1
Gda
Dlk1
Drd3

**Supplementary Figure 2: Examples of *in situ* hybridization for differentially expressed genes in PFC DS and NAc.**

*In situ* hybridization patterns from the Allen Brain Institute (http://mouse.brain-map.org/) for selected genes that were also studied by RT-PCR (see **Fig. 3b-d**). Images are shown for genes that were found by BAC-TRAP/RNA-Seq and QRT-PCR to be preferentially expressed in PFC (**a**), in DS (**b**), or in NAc (**c**). For gene products with strong enrichment and high expression levels the hybridization differences are easily visible (e.g. *Tbr1* in cortex, *Slc24a2* in DS, or *Wfs1*, *Gda* or *Drd3* in NAc), whereas for other gene products, RNA-Seq and RT-PCR are more informative.

**a**

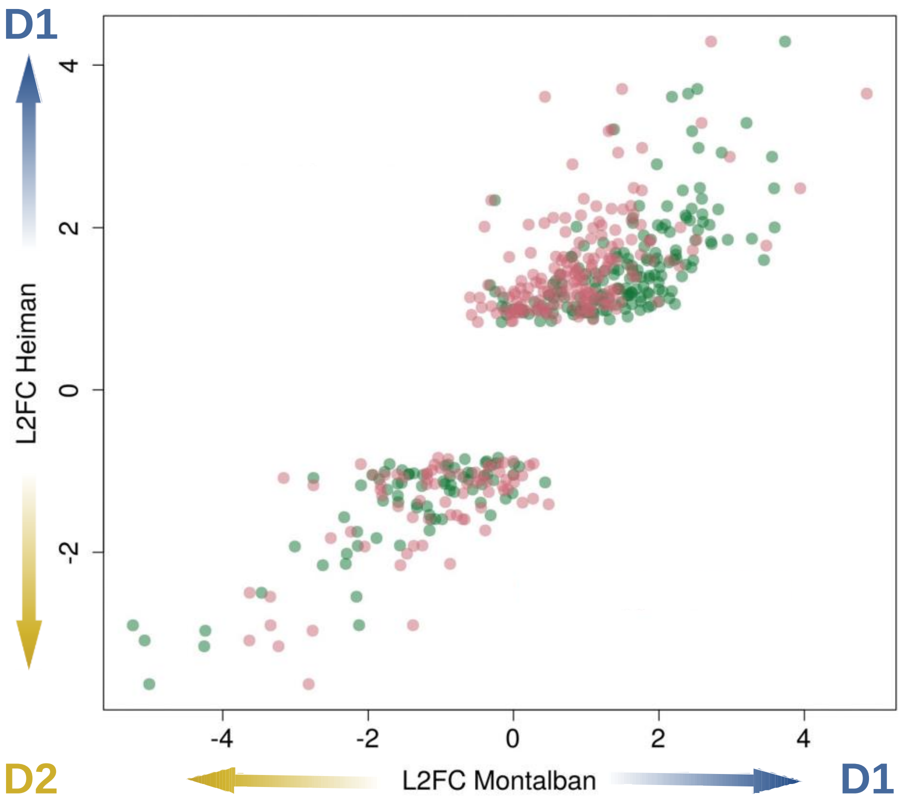

**a**

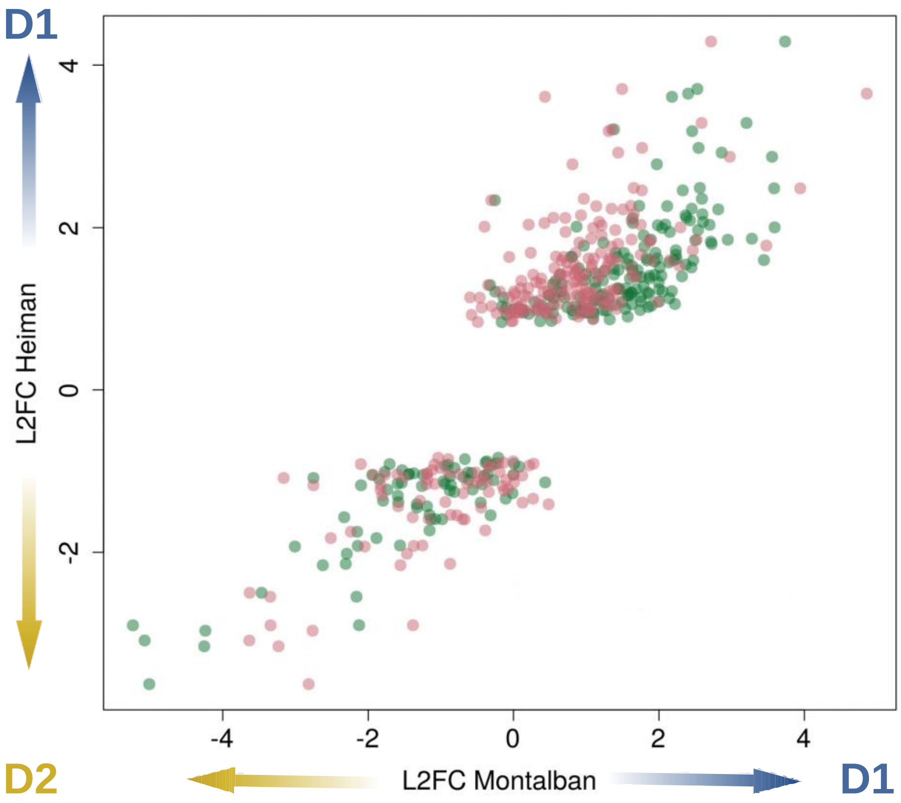

c

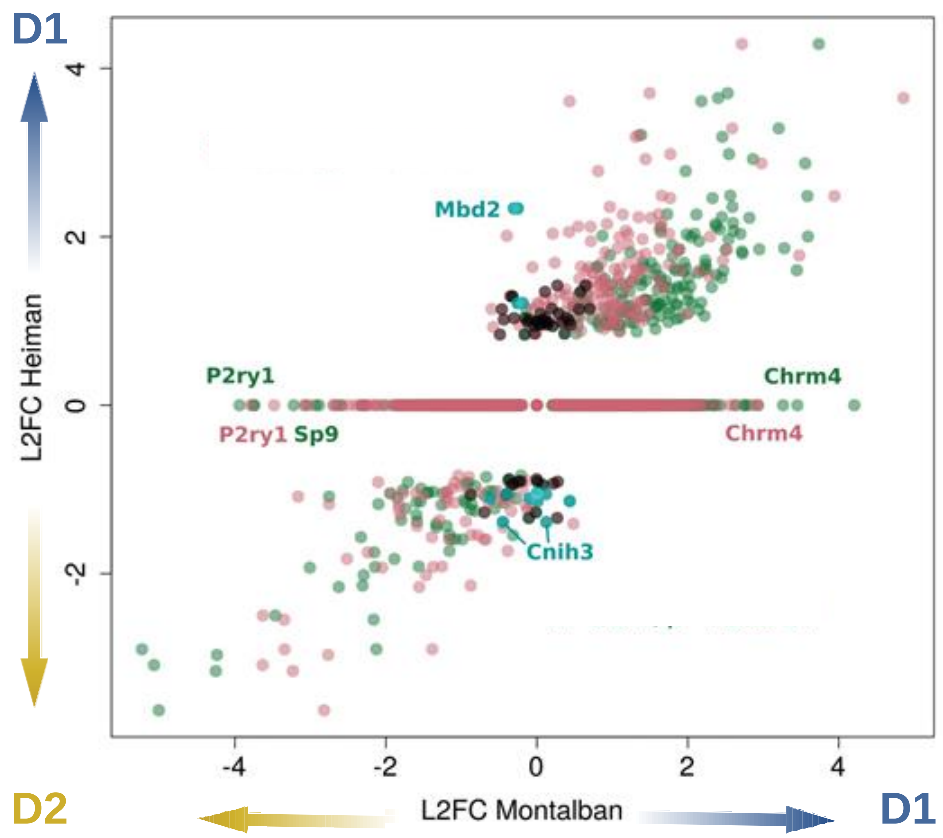
D1
D1
D2
L2FC Heiman
L2FC Montalban
4
2
0
-2
-4
Mbd2
P2ry1
Chrm4
P2ry1
Sp9
Chrm4
Cnih3

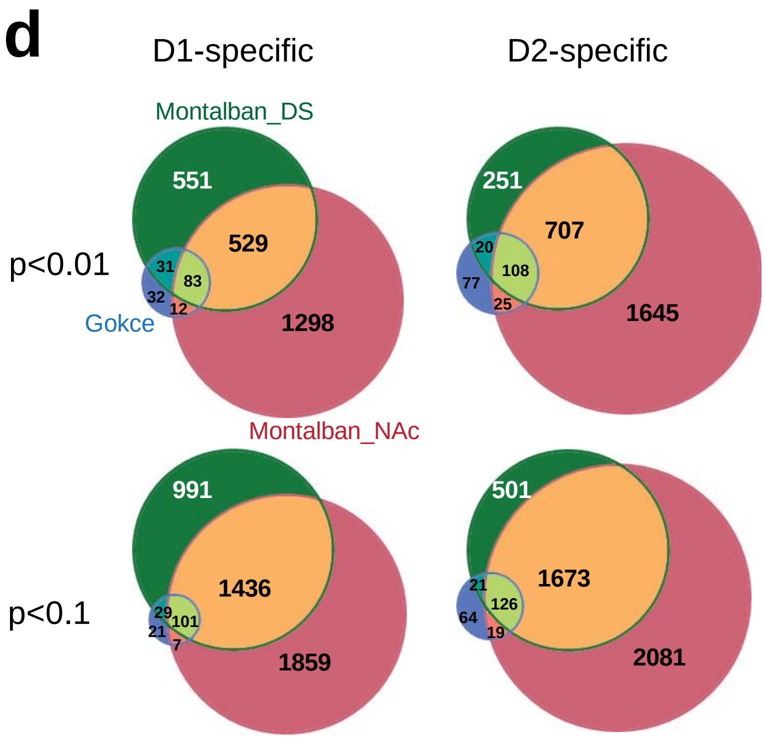
d
D1-specific
D2-specific
Montalban_DS
551
529
31
83
32
12
Gokce
1298
251
707
20
108
77
25
1645
p<0.01
Montalban_NAc
991
1436
29
101
21
7
1859
501
1673
21
126
64
19
2081
p<0.1

**Supplementary Figure 3: Comparison of the results of the current study with previous reports on D1 D2 differences.**

**a**. mRNAs differentially expressed between D1 and D2 neurons in Heiman et al.[1] (Y axis, log2 fold-changes [L2FC] Heiman) and in the current study (X axis, L2FC Montalban) in the DS (green) or the NAc (magenta). After exclusion of the BAC genes, 248 mRNAs were confirmed in the current study and the correlation between L2FC was very good (DS, R = 0.91, p = 4.9 $10^{-94}$, NAc, R = 0.83, p = 1.54 $10^{-63}$). Note that each gene product was plotted twice, against our values in DS and NAc. **b.** Same as in **a.** but including also the 33 mRNAs found to be differentially expressed in Heiman et al. but not here (black). With the exception of Mbd2, the L2FC changes observed for those mRNAs are on the lower end of the scale in both studies. The 5042 mRNAs found differentially expressed in our study but not in Heiman et al. are also shown (dots aligned horizontally on L2FC Heiman 0). They include mRNAs whose expression differs by up to one order of magnitude between D1 and D2 neurons (horizontal dispersion of the dots at y = 0; e.g. *P2ry1*, *Sp9* both enriched in D2 neurons and *Chrm4*, enriched in D1 neurons). **c.** Same as in **b**. but also indicating 7 mRNAs (blue) identified as different between D1 and D2 by Heiman et al. but neither in our current study nor in Ho et al.[2]. They include *Mdb2* and the cornichon 3 homolog *Cnih3*, which showed the strongest absolute L2FC in Heiman *et* al. Excluding these mRNAs reduces the range of log2 fold changes for our possible false negatives to [-1.34, 1.42]. **d.** Venn diagram comparing the results of our study with those of Gokce *et al*.[3]. Eighty % of their D1-specific mRNAs were also found D1-specific in our study, as were 67 % of their D2-specific mRNAs. Most of the genes we did not confirm, exhibited a low expression (e.g. *Rbp4*) and/or a low fold-change (|L2FC|<1), and often an opposite D1 vs D2 specificity (**Supplementary Tables 14a-b**). Increasing the adjusted p-value significance threshold from <0.01 to <0.1 (as in Ref[1]) in our study, recovered only 11 and 13 genes specific for D1 and D2 respectively, found differentially expressed in Refs[1] and[2], but not in another single cell RNA-Seq analysis[3]. Conversely, using such a less stringent threshold would have increased the number of mRNAs differentially expressed between D1 and D2 neurons to 4878 in the DS and 7302 in the NAc, probably containing a large number of false positive.

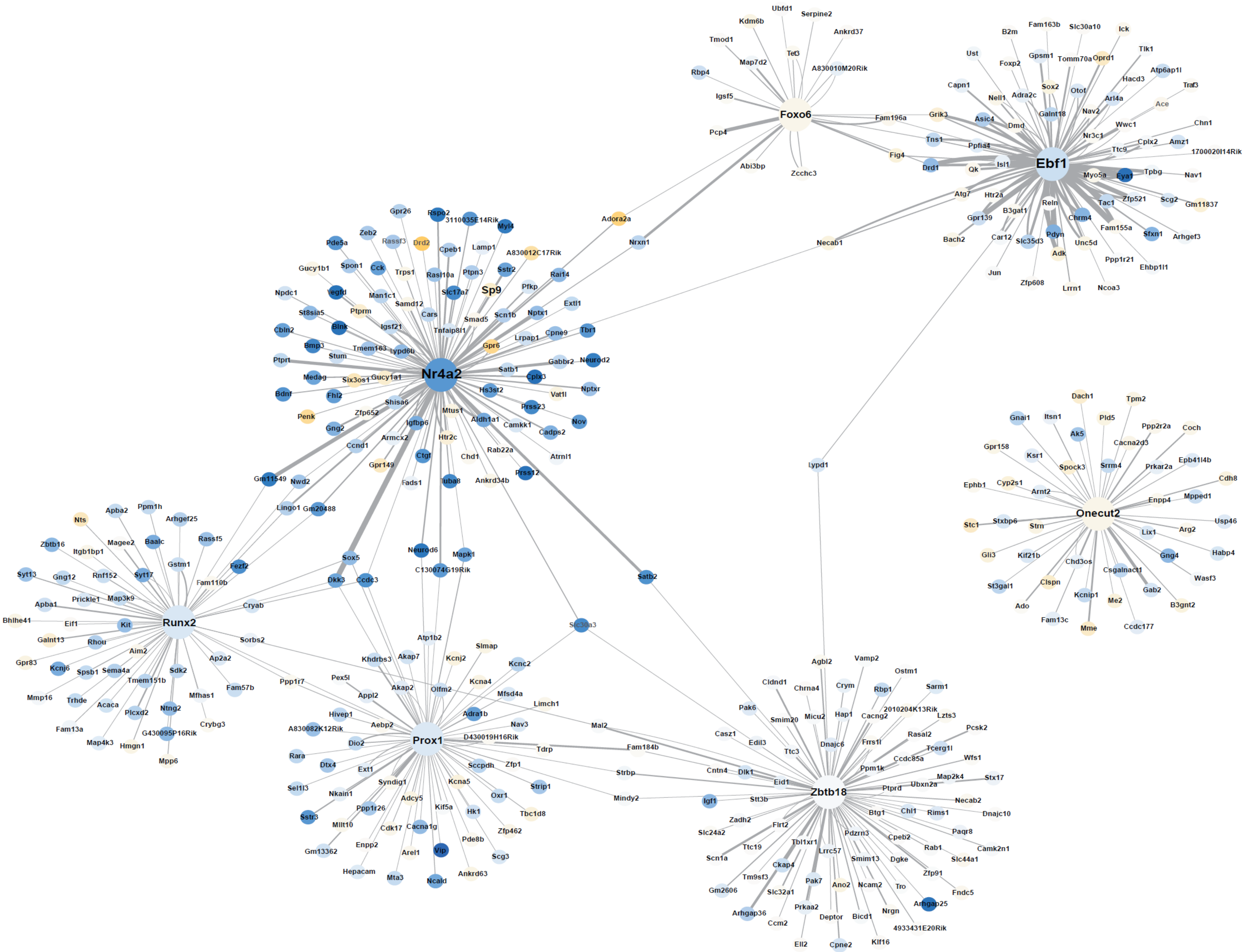

Ubfd1
Kdm6b
Serpine2
Ankrd37
Tmod1
Tet3
Map7d2
A830010M20Rik
Rbp4
Igsf5
Foxo6
Fam196a
Pcp4
Abi3bp
Zcchc3
Fam163b
Slc30a10
Ick
B2m
Tlk1
Ust
Gpsm1
Tomm70a
Oprd1
Foxp2
Atp6ap1l
Capn1
Hacd3
Traf3
Sox2
Otof
Nell1
Adra2c
Arl4a
Ace
Galnt18
Grik3
Nav2
Asic4
Dmd
Wwc1
Chn1
Nr3c1
Tns1
Ppfia4
Cplx2
Amz1
Ttc9
1700020I14Rik
Fig4
Isl1
Drd1
Qk
Ebf1
Myo5a
Eya1
Tpbg
Nav1
Atg7
Htr2a
Zfp521
Scg2
Gm11837
Tac1
Reln
B3gat1
Gpr139
Chrm4
Fam155a
Sfxn1
Arhgef3
Bach2
Car12
Slc35d3
Pdyn
Unc5d
Adk
Ppp1r21
Ehbp1l1
Jun
Zfp608
Lrrn1
Ncoa3
Necab1
Adora2a
Nrxn1
Gpr26
Rspo2
3110035E14Rik
Myl4
Zeb2
Pde5a
Rassf3
Drd2
Cpeb1
Lamp1
A830012C17Rik
Gucy1b1
Spon1
Cck
Trps1
Rasl10a
Ptpn3
Sstr2
Rai14
Npdc1
Vegfd
Man1c1
Slc17a7
Sp9
Pfkp
Samd12
Extl1
St8sia5
Ptprm
Cars
Smad5
Scn1b
Nptx1
Cbln2
Blnk
Igsf21
Tnfaip8l1
Lrpap1
Cpne9
Tbr1
Bmp3
Tmem163
Lypd6b
Gpr6
Stum
Ptprt
Gabbr2
Neurod2
Medag
Six3os1
Gucy1a1
Nr4a2
Satb1
Cplx3
Bdnf
Fhl2
Hs3st2
Vat1l
Nptxr
Shisa6
Penk
Zfp652
Mtus1
Prss23
Aldh1a1
Camkk1
Nov
Igfbp6
Gng2
Cadps2
Htr2c
Ccnd1
Armcx2
Rab22a
Ctgf
Chd1
Atrnl1
Gpr149
Gm11549
Nwd2
Fads1
Tuba8
Ankrd34b
Prss12
Lingo1
Gm20488
Ppm1h
Apba2
Nts
Arhgef25
Zbtb16
Magee2
Baalc
Rassf5
Itgb1bp1
Gstm1
Fezf2
Syt13
Gng12
Rnf152
Syt17
Fam110b
Prickle1
Map3k9
Apba1
Cryab
Bhlhe41
Eif1
Kit
Runx2
Galnt13
Rhou
Sorbs2
Aim2
Gpr83
Kcnj6
Spsb1
Sema4a
Sdk2
Ap2a2
Tmem151b
Mmp16
Trhde
Acaca
Mfhas1
Fam57b
Ppp1r7
Fam13a
Plcxd2
Ntng2
G430095P16Rik
Crybg3
Map4k3
Hmgn1
Mpp6
Sox5
Dkk3
Ccdc3
Neurod6
Mapk1
C130074G19Rik
Satb2
Slc30a3
Lypd1
Atp1b2
Slmap
Khdrbs3
Akap7
Kcnj2
Kcnc2
Pex5l
Kcna4
Akap2
Olfm2
Mfsd4a
Appl2
Limch1
Hivep1
Adra1b
Nav3
Mal2
Aebp2
A830082K12Rik
Prox1
D430019H16Rik
Dio2
Tdrp
Fam184b
Rara
Dtx4
Ext1
Sccpdh
Zfp1
Syndig1
Kcna5
Sel1l3
Strip1
Strbp
Nkain1
Adcy5
Oxr1
Mindy2
Kif5a
Ppp1r26
Sstr3
Hk1
Tbc1d8
Mllt10
Cdk17
Cacna1g
Zfp462
Pde8b
Gm13362
Enpp2
Arel1
Vip
Scg3
Hepacam
Mta3
Ncald
Ankrd63
Dach1
Tpm2
Gnai1
Itsn1
Pld5
Ppp2r2a
Coch
Ak5
Cacna2d3
Gpr158
Ksr1
Srrm4
Prkar2a
Epb41l4b
Spock3
Cdh8
Ephb1
Cyp2s1
Arnt2
Enpp4
Mpped1
Onecut2
Stc1
Stxbp6
Strn
Lix1
Arg2
Usp46
Gng4
Habp4
Gli3
Kif21b
Chd3os
Csgalnact1
Wasf3
St3gal1
Clspn
Gab2
Kcnip1
Me2
Ado
B3gnt2
Fam13c
Mme
Ccdc177
Agbl2
Vamp2
Ostm1
Cldnd1
Chrna4
Crym
Rbp1
Sarm1
Pak6
2010204K13Rik
Smim20
Micu2
Hap1
Cacng2
Lzts3
Pcsk2
Casz1
Edil3
Rasal2
Dnajc6
Frrs1l
Ttc3
Tcerg1l
Ccdc85a
Wfs1
Cntn4
Dlk1
Ppm1k
Map2k4
Stx17
Eid1
Ptprd
Ubxn2a
Igf1
Stt3b
Zbtb18
Necab2
Btg1
Chl1
Rims1
Dnajc10
Zadh2
Flrt2
Slc24a2
Pdzrn3
Paqr8
Cpeb2
Tbl1xr1
Rab1
Camk2n1
Ttc19
Lrrc57
Scn1a
Smim13
Dgke
Slc44a1
Ckap4
Zfp91
Tm9sf3
Pak7
Ano2
Ncam2
Tro
Fndc5
Gm2606
Slc32a1
Arhgap25
Arhgap36
Prkaa2
Nrgn
Deptor
Bicd1
Ccm2
4933431E20Rik
Ell2
Cpne2
Kif16

**Supplementary Figure 4: Striatal gene network of transcription factors illustrating connections with mRNAs differentially expressed in D1 and D2 cells.**

The network was generated using an expression-based network inference procedure and combined the results of CLR[4], a mutual-information-based approach providing undirected edges, and GENIE3[5], a tree-based regression approach providing directed edges, using filtered CPM (see **Online Methods**). Subnetworks were extracted using gene lists as seeds, retaining only the first neighbors with scores above a threshold. Visualization and analysis of the resulting networks was done using Cytoscape[6], only keeping the 1 % edges with the highest scores. The edges that connected either one or two of the TFs differentially expressed between DS and NAc focusing on TFs specific of D1 cells are shown. Seven TFs passed the filter and six of them formed a connected subnetwork of clusters (i.e. sharing first neighbors). Nodes are colored according to their relative expression in D1 (blue) and D2 (yellow) neurons. *Nr4a2* and *Ebf1* are surrounded by genes differentially expressed in D1 and D2 cells. The color intensities reflect differential expression and line thickness the reliability of prediction.

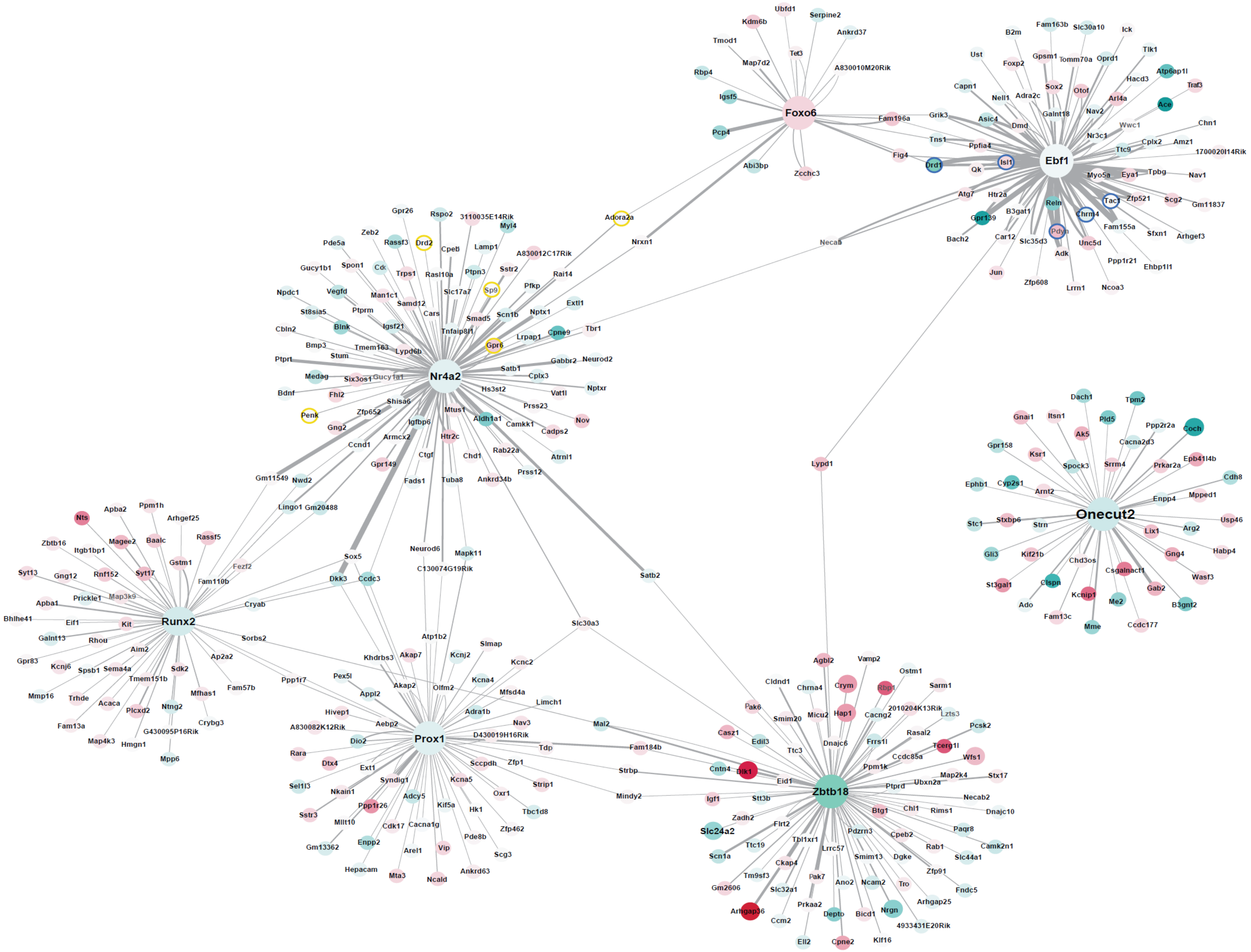
Foxo6
Ubfd1
Serpine2
Kdm6b
Ankrd37
Tmod1
Tet3
Map7d2
A830010M20Rik
Rbp4
Igsf5
Pcp4
Abi3bp
Zcchc3
Fam196a
Fig4
Ebf1
B2m
Fam163b
Slc30a10
Ick
Tlk1
Ust
Gpsm1
Tomm70a
Oprd1
Foxp2
Hacd3
Atp6ap1l
Capn1
Sox2
Otof
Traf3
Nell1
Adra2c
Arl4a
Ace
Galnt18
Nav2
Grik3
Asic4
Dmd
Wwc1
Chn1
Nr3c1
Tns1
Ppfia4
Ttc9
Cplx2
Amz1
1700020I14Rik
Drd1
Qk
Isl1
Myo5a
Eya1
Tpbg
Nav1
Atg7
Htr2a
Tac1
Zfp521
Scg2
Gm11837
B3gat1
Reln
Gpr139
Chrm4
Fam155a
Sfxn1
Arhgef3
Bach2
Car12
Slc35d3
Pdyn
Unc5d
Adk
Ppp1r21
Ehbp1l1
Jun
Zfp608
Lrrn1
Ncoa3
Necab
Nr4a2
Gpr26
Rspo2
3110035E14Rik
Myl4
Zeb2
Pde5a
Rassf3
Drd2
Cpeb
Lamp1
A830012C17Rik
Gucy1b1
Spon1
Cd
Trps1
Rasl10a
Ptpn3
Sstr2
Rai14
Npdc1
Vegfd
Man1c1
Slc17a7
Sp9
Pfkp
Samd12
St8sia5
Ptprm
Cars
Smad5
Scn1b
Nptx1
Extl1
Igsf21
Blnk
Cbln2
Tnfaip8l1
Lrpap1
Cpne9
Tbr1
Bmp3
Tmem163
Gpr6
Lypd6b
Stum
Ptprt
Gabbr2
Neurod2
Medag
Six3os1
Gucy1a1
Satb1
Cplx3
Bdnf
Fhl2
Hs3st2
Vat1l
Nptxr
Shisa6
Prss23
Penk
Zfp652
Mtus1
Nov
Igfbp6
Aldh1a1
Camkk1
Cadps2
Gng2
Armcx2
Htr2c
Ccnd1
Rab22a
Atrnl1
Ctgf
Chd1
Gpr149
Gm11549
Nwd2
Fads1
Tuba8
Ankrd34b
Prss12
Lingo1
Gm20488
Adora2a
Nrxn1
Runx2
Nts
Apba2
Ppm1h
Arhgef25
Zbtb16
Magee2
Baalc
Rassf5
Itgb1bp1
Gstm1
Fezf2
Syt13
Gng12
Rnf152
Syt17
Fam110b
Prickle1
Map3k9
Apba1
Bhlhe41
Eif1
Kit
Cryab
Galnt13
Rhou
Sorbs2
Aim2
Ap2a2
Gpr83
Kcnj6
Spsb1
Sema4a
Sdk2
Tmem151b
Fam57b
Mmp16
Trhde
Mfhas1
Acaca
Ntng2
Plcxd2
Fam13a
Crybg3
G430095P16Rik
Map4k3
Hmgn1
Mpp6
Sox5
Dkk3
Ccdc3
Neurod6
Mapk11
C130074G19Rik
Satb2
Slc30a3
Prox1
Atp1b2
Slmap
Khdrbs3
Akap7
Kcnj2
Kcnc2
Ppp1r7
Pex5l
Akap2
Kcna4
Appl2
Olfm2
Mfsd4a
Limch1
Hivep1
Adra1b
A830082K12Rik
Aebp2
Nav3
Mal2
Dio2
D430019H16Rik
Rara
Tdp
Fam184b
Dtx4
Sccpdh
Zfp1
Ext1
Strbp
Sel1l3
Syndig1
Strip1
Kcna5
Nkain1
Adcy5
Oxr1
Mindy2
Sstr3
Ppp1r26
Kif5a
Hk1
Tbc1d8
Mllt10
Cdk17
Cacna1g
Zfp462
Gm13362
Enpp2
Arel1
Vip
Pde8b
Scg3
Hepacam
Mta3
Ncald
Ankrd63
Onecut2
Dach1
Tpm2
Gnai1
Itsn1
Pld5
Ppp2r2a
Coch
Ak5
Gpr158
Cacna2d3
Ksr1
Spock3
Srrm4
Prkar2a
Epb41l4b
Ephb1
Cyp2s1
Cdh8
Arnt2
Enpp4
Mpped1
Stc1
Stxbp6
Strn
Usp46
Lix1
Arg2
Gli3
Kif21b
Gng4
Habp4
Chd3os
Csgalnact1
Wasf3
St3gal1
Clspn
Gab2
Kcnip1
Me2
B3gnt2
Ado
Fam13c
Mme
Ccdc177
Lypd1
Zbtb18
Agbl2
Vamp2
Ostm1
Cldnd1
Chrna4
Crym
Rbp1
Sarm1
Pak6
Smim20
Micu2
Hap1
Cacng2
2010204K13Rik
Lzts3
Pcsk2
Casz1
Edil3
Dnajc6
Frrs1l
Rasal2
Ttc3
Tcerg1l
Wfs1
Ccdc85a
Cntn4
Dlk1
Ppm1k
Map2k4
Stx17
Eid1
Ptprd
Ubxn2a
Igf1
Stt3b
Necab2
Btg1
Chl1
Rims1
Dnajc10
Zadh2
Slc24a2
Flrt2
Pdzrn3
Paqr8
Cpeb2
Tbl1xr1
Rab1
Camk2n1
Ttc19
Lrrc57
Scn1a
Smim13
Dgke
Slc44a1
Ckap4
Tm9sf3
Zfp91
Pak7
Gm2606
Ano2
Ncam2
Tro
Fndc5
Slc32a1
Arhgap25
Prkaa2
Arhgap36
Nrgn
Depto
Bicd1
Ccm2
4933431E20Rik
Ell2
Cpne2
Klf16

**Supplementary Fig. 5: Striatal gene network of transcription factors illustrating connections with genes differentially expressed in DS and NAc.**

Same network as in **Supplementary Figure 5**, except that nodes are colored according to their relative expression in DS (green) and NAc (red). Genes linked to *Onecut2* and *Zbtb18* are strongly differentially expressed in DS and NAc. The color intensities reflect differential expression and line thickness the reliability of prediction. For information, the genes connected to *Nr4a2* and *Ebf1* and typical of D1 or D2 cells (see Supplementary Figure 5) are indicated by a blue or yellow lining respectively.

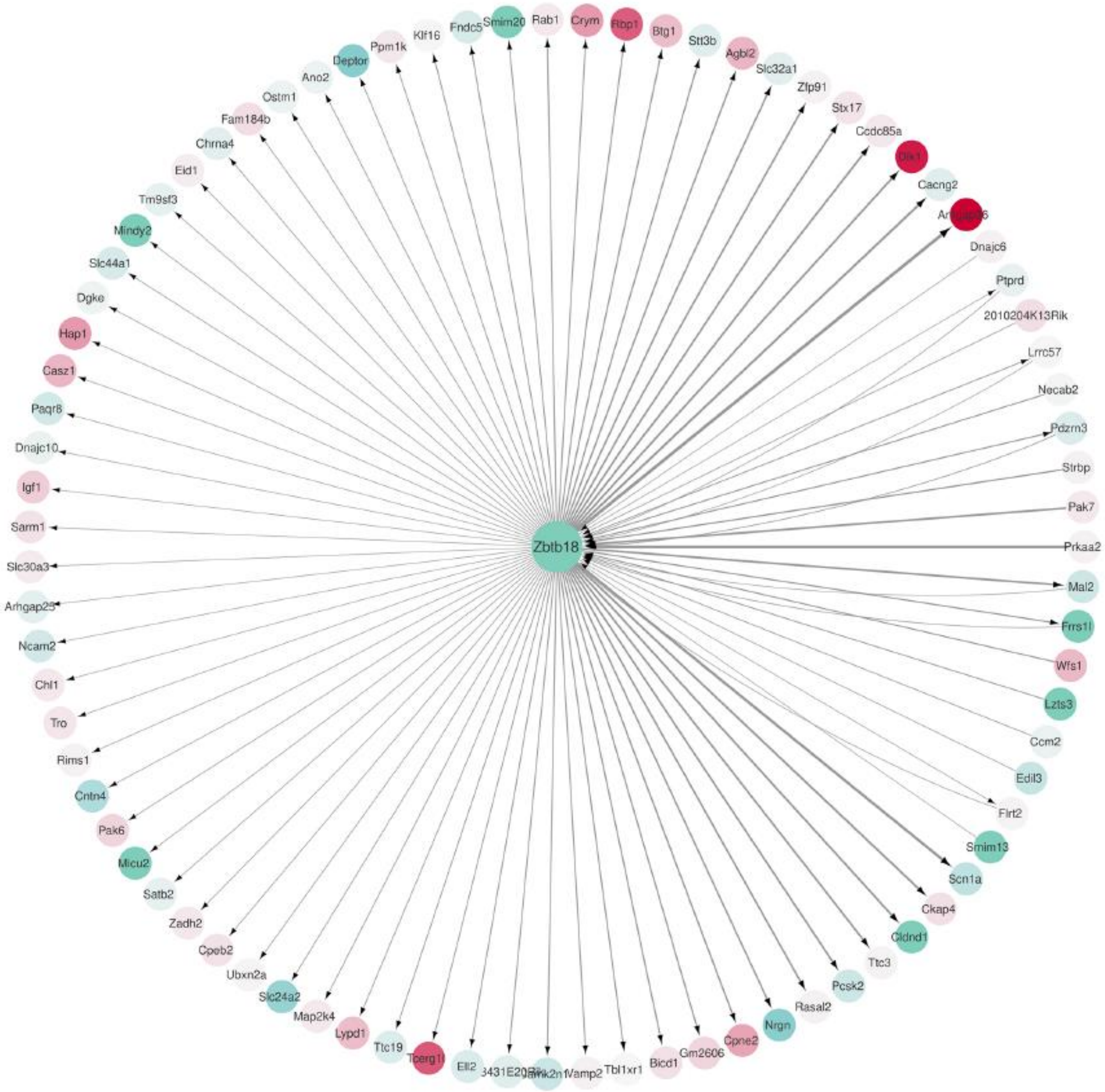


**Supplementary Figure 6: Gene expression-based network inference of Zbtb18 regulations in SPNs.** Only influences passing our threshold are shown. Arrows indicate the direction of the influences. Line thickness reflects the reliability of prediction. Gene nodes are colored according to their relative expression in DS (green) or NAc (red). Among threshold-passing influences, 66 are coming solely from *Zbtb18*, 10 are solely going to *Zbtb18* and 7 are bidirectional, but with a higher weight of outgoing interaction.

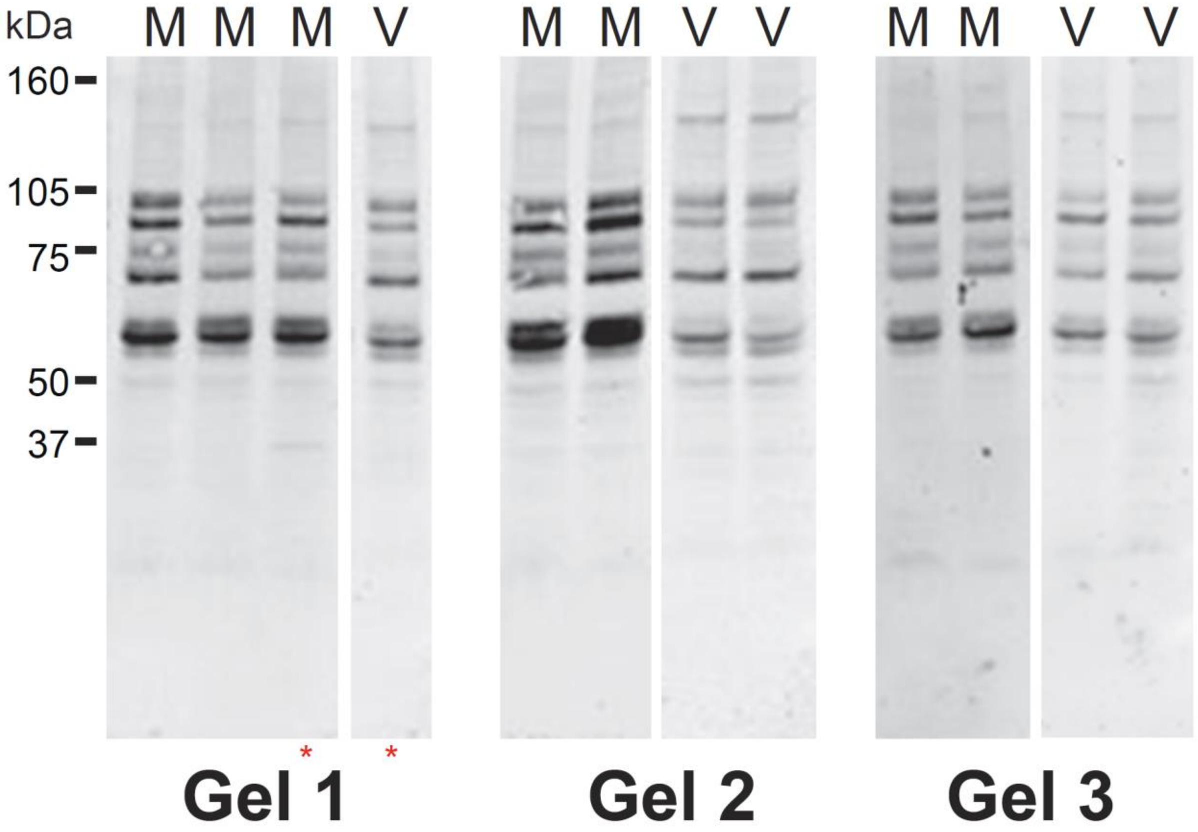


**Supplementary Fig. 7: Immunoblots of pPKA-substrates quantified in Fig. 6a.**

Wild type mice were treated with vehicle (V) or misoprostol (M, 0.1 mg.kg$^{-1}$ i.p.) 30 min before sacrifice and DS homogenates analyzed by immunoblotting with anti-phospho-PKA-substrate. Three different gels were run, as indicated, from which the relevant lanes are shown. The positions of molecular weight markers are indicated. *, lanes shown in **Fig. 6a**.

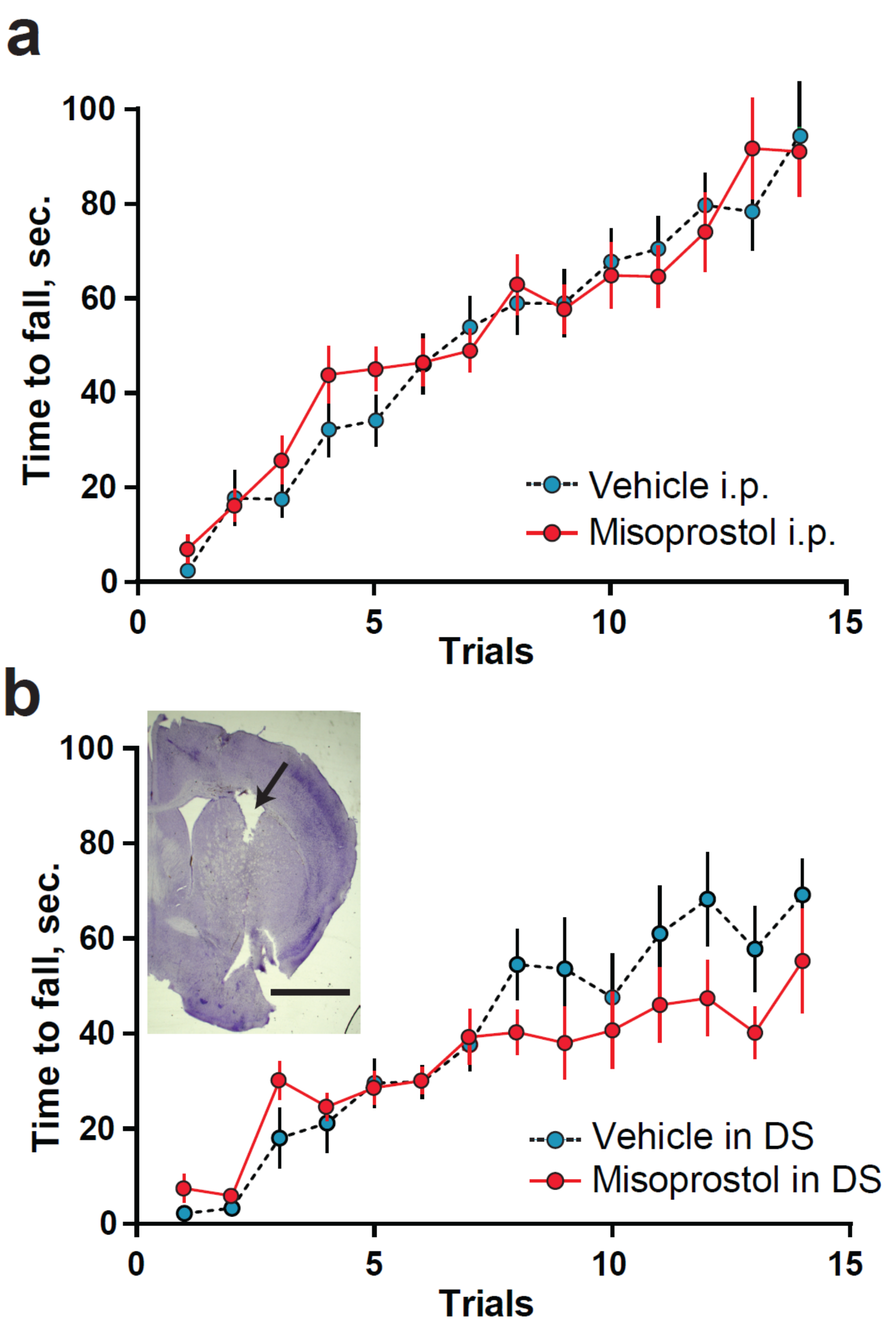


**Supplementary Fig. 8: Lack of effect of a prostaglandin agonist on rotarod learning.**

**a.** Wild type mice were implanted on day 1 with an i.p. osmotic minipump containing either vehicle (Veh) or misoprostol (Miso, 50 µg $kg^{-1}$ $day^{-1}$). They were tested on accelerating rotarod starting at day 9 (4 trials per day). Latency to fall during the learning phase was recorded. Two-way repeated measures ANOVA showed an effect of time ($F_{(11,\ 562)}$ = 23.36, p<0.0001), but not of treatment ($F_{(11,562)}$ = 0.05, p = 0.81), n = 14 mice per group (detailed statistical results in **Supplementary Table 19**). **b.** Wild type mice were implanted on day 1 with two subcutaneous osmotic minipumps containing Veh or Miso and linked to a catheter connected to a cannula implanted in each DS, as in **Fig. 6b**. Mice were tested in accelerating rotarod as in **a**. Two-way repeated measures ANOVA showed an effect of time ($F_{(13,\ 221)}$ = 28.68, p<0.0001), but not of treatment ($F_{(1,221)}$ = 0.64, p = 0.43), n = 9-10 mice per group (detailed statistical results in **Supplementary Table 19**). **Inset:** Nissl-stained coronal brain section showing a cannula track (**arrow**) in the DS. Scale bar, 2 mm.
