## Supplementary material for "Translational profiling of mouse dopaminoceptive neurons reveals a role of PGE2 in dorsal striatum": List of supplementary Tables

Enrica Montalban et al.

### LIST OF SUPPLEMENTARY TABLES

**Supplementary Table 1: All_data.**

1a- Sample description incl. sex of mice
1b- Library description
1c- Count Table
1d- Cell type markers
1e- References for Table 1d

**Supplementary Table 2: Complete-DESeq-results.**

**Supplementary Table 3: DE-genes_PFC-vs-STR.**

Analytical comparisons of differentially expressed genes between PFC and STR (i.e. DS + NAc), all in D1 samples (data from Supplementary Table 2).

3a- DE genes PFC>Str
3b- DE genes Str>PFC
3c- DE genes PFC>Str best (padj<0.001, L2FC>2)
3d- DE genes Str>PFC best (padj<0.001, L2FC>2)
3e- All PFC samples > all Str samples
3f- All Str samples > all PFC samples
3g- GO analysis mRNAs PFC>Str
3h- GO analysis mRNAs Str>PFC
3i- IUAPHAR-identified mRNAs PFC>Str
3j- IUAPHAR-identified mRNAs Str>PFC

**Supplementary Table 4: DEXSeq_PFC-vs-Str_in-D1**.

Differentially expressed exons (DEXSeq) in PFC vs Str in D1 samples.

Exon usage analysis by the DEXseq package. DEXseq breaks down open reading frames into exon fragments (which do not necessarily correspond to full exons), and tests if there is a significant change in the relative usage of each of these fragments, i.e. in the ratio of sequencing reads mapping to this fragment, over the reads mapping to any other fragment of the same gene. FeatureID corresponds to the arbitrary numbering of the exonic fragment by DEXseq, and the genomic coordinates for each fragment are given in the genomicData.seqnames, genomicData.start and genomicData.end columns. padj is the multiple testing correction adjusted p-value for the differential usage of a given fragment, and log2fold corresponds to the fold-change of usage for that fragment.

**Supplementary Table 5: DEXSeq_PFC-vs-Str_in-D1_Analyses.**
Analyses of results in Supplementary Table 4 *(see legend to Supplementary Table 4).*

5a- Exons PFC>Str padj<0.05
5b- Exons Str>PFC padj<0.05
5c- Gene counts
5d- Genes with at least one exon PFC>Str (Table 5a) with increased total mRNA (Table 3a)
5e- Genes with at least one exon Str>PFC (Table 5b) with increased total mRNA (Table 3b)
5f- *Arpp21* exon usage PFC vs Str

**Supplementary Table 6: DE-genes_D1-vs-D2_in-Str.**
Analytical comparisons of differentially expressed genes between D1 and D2 in DS and NAc samples (data from Supplementary Table 2).

6a- DE genes D1>D2 in both DS and NAc
6b- DE genes D1>D2 only in DS
6c- DE genes D1>D2 only in NAc
6d- DE genes D2>D1 in both DS and NAc
6e- DE genes D2>D1 only in DS
6f- DE genes D2>D1 only in NAc
6g- DE genes with opposite variation in DS and NAc
6h- Best DE genes D1>D2 in DS (padj<0.01, L2FC>1)
6i- Best DE genes D1>D2 in NAc (padj<0.01, L2FC>1)
6j- Best DE genes D1>D2 common between DS and NAc
6k- Best DE genes D2>D1 in DS (padj<0.01, L2FC>1)
6l- Best DE genes D2>D1 in NAc (padj<0.01, L2FC>1)
6m- Best DE genes D2>D1 common between DS and NAc
6n- All D1 samples > all D2 samples in DS
6o- All D1 samples > all D2 samples in NAc
6p- All D1 samples > all D2 samples in DS and NAc (common to tables 6n and 6o)
6q- All D2 samples > all D1 samples in DS
6r- All D2 samples > all D1 samples in NAc
6s- All D2 samples > all D1 samples in DS and NAc (common to tables 6q and 6r)

**Supplementary Table 7: DE-genes_D1-vs-D2_in-STR_GO-IUAPHAR.**
Analyses of mRNAs differentially expressed in D1 vs D2 samples (Supplementary Tables 6a-g) using gene ontology and IUAPHAR classification.

7a- GO analysis mRNAs D1>D2 in both DS and NAc
7b- GO analysis mRNAs D1>D2 only in DS
7c- GO analysis mRNAs D1>D2 only in NAc
7d- GO analysis mRNAs D2>D1 in both DS and NAc
7e- GO analysis mRNAs D2>D1 only in DS
7f- GO analysis mRNAs D2>D1 only in NAc
7g- IUAPHAR-identified mRNAs D1>D2
7h- IUAPHAR-identified mRNAs D2>D1

**Supplementary Table 8: DEXSeq_D1-vs-D2_in-DS.**
Differentially expressed exons (DEXSeq) between D1 and D2 in DS samples *(see legend to Supplementary Table 4).*

**Supplementary Table 9: DEXSeq_D1-vs-D2_in-NAc**.
Differentially expressed exons (DEXSeq) between D1 and D2 in NAc samples *(see legend to Supplementary Table 4).*

**Supplementary Table 10: DEXSeq_D1-vs-D2_in-DS-and-NAc_Analyses.**
Analyses of DEXSeq results in DS (Supplementary Table 8) and NAc (Supplementary Table 9), *and see legend to Supplementary Table 4.*

10a- Exons D1>D2 in DS padj<0.05
10b- Exons D2>D1 in DS padj<0.05
10c- Exons D1>D2 in NAc padj<0.05
10d- Exons D2>D1 in NAc padj<0.05
10e- Gene counts
10f- Genes with at least one exon D1>D2 in DS (Table 10a) with increased total mRNA (Tables 7a,b)
10g- Genes with at least one exon D2>D1 in DS (Table 10b) with increased total mRNA (Tables 7d,e)
10h- Genes with at least one exon D1>D2 in NAc (Table 10c) with increased total mRNA (Tables 7a,c)
10i- Genes with at least one exon D2>D1 in NAc (Table 10d) with increased total mRNA (Tables 7d,f)

**Supplementary Table 11: DE-genes_DS-vs-NAc.**
Analytical comparisons of differentially expressed genes between DS and NAc in D1 and D2 samples (data from Supplementary Table 2).

11a- DE genes DS>NAc in both D1 and D2
11b- DE genes DS>NAc only in D1
11c- DE genes DS>NAc only in D2
11d- DE genes NAc>DS in both D1 and D2
11e- DE genes NAc>DS only in D1
11f- DE genes NAc>DS only in D2
11g- DE genes opposite variation in D1 and D2
11h- Best DE genes DS>NAc in D1 (padj<0.01, L2FC>1)
11i- Best DE genes DS>NAc in D2 (padj<0.01, L2FC>1)
11j- Best DE genes DS>NAc common between D1 and D2
11k- Best DE genes NAc>DS in D1 (padj<0.01, L2FC>1)
11l- Best DE genes NAc>DS in D2 (padj<0.01, L2FC>1)
11m- Best DE genes NAc>DS common between D1 and D2
11n- All DS samples > all NAc samples in D1
11o- All DS samples > all NAc samples in D2
11p- All DS samples > all NAc samples in D1 and D2 (common to tables 11n and 11o)
11q- All NAc samples > all DS samples in D1
11r- All NAc samples > all DS samples in D2
11s- All NAc samples > all DS samples in D1 and D2 (common to tables 11q and 11r)
11t- Selection of top differentially expressed genes between DS and NAc (most abundant and/or most differentially expressed)

**Supplementary Table 12: DE-genes_DS-vs-NAc_GO-IUAPHAR.**
Analyses of mRNAs differentially expressed in DS vs NAc (Supplementary Tables 11a-g) using gene ontology and IUAPHAR classification.

12a- GO analysis mRNAs DS>NAc in both D1 and D2
12b- GO analysis mRNAs DS>NAc only in D1

12c- GO analysis mRNAs DS>NAc only in D2
12d- GO analysis mRNAs NAc>DS in both D1 and D2
12e- GO analysis mRNAs NAc>DS only in D1
12f- GO analysis mRNAs NAc>DS only in D2
12g- IUAPHAR-identified mRNAs DS>NAc
12h- IUAPHAR-identified mRNAs NAc>DS

**Supplementary Table 13: DEXSeq_DS-vs-NAc_in-D1.**
Differentially expressed exons (DEXSeq) between DS and NAc in D1 samples *(see legend to Supplementary Table 4)*.

**Supplementary Table 14: DEXSeq_DS-vs-NAc_in-D2.**
Differentially expressed exons (DEXSeq) between DS and NAc in D2 samples *(see legend to Supplementary Table 4)*.

**Supplementary Table 15: DEXSeq_DS-vs-NAc_in-D1-and-D2_Analyses.**
Analyses of DEXSeq results DS vs NAc, in D1 (Supplementary Table 13) and NAc (Supplementary Table 14) samples *(see legend to Supplementary Table 4)*.

15a- Exons DS>NAc in D1 padj<0.05
15b- Exons NAc>DS in D1 padj<0.05
15c- Exons DS>NAc in D2 padj<0.05
15d- Exons NAc>DS in D2 padj<0.05
15e- Gene counts
15f- Genes with at least one exon DS>NAc in D1 (Table 15a) with increased total mRNA (Table 12a,d)
15g- Genes with at least one exon NAc>DS in D1 (Table 15b) with increased total mRNA (Table 12b,e)
15h- Genes with at least one exon DS>NAc in D2 (Table 15c) with increased total mRNA (Table 12a,f)
15i- Genes with at least one exon NAc>DS in D2 (Table 15d) with increased total mRNA (Table 15b,g)

**Supplementary Table 16: DE-Transcription-factors_STR.**
Differentially expressed transcription factors (TF) in the striatum (data from Supplementary Tables 6 and 11).

16a- DE TFs D1 vs D2 common to DS and NAc
16b- DE TFs D1 vs D2 DS-specific
16c- DE TFs D1 vs D2 NAc-specific
16d- DE TFs DS vs NAc common to D1 and D2
16e- DE TFs DS vs NAc D1-specific
16f- DE TFs DS vs NAc D2-specific

**Supplementary Table 17: IPA-upstream-analysis_DS-vs-NAc.**
Ingenuity pathway analysis (IPA-Partek) of DS vs NAc differences combining D1 and D2 neurons (limited to potentially endogenous factors).

**Supplementary Table 18: Prostaglandin-related-gene-products.**
Expression of prostaglandin-related genes in striatal samples (data from Supplementary Table 2).

**Supplementary Table 19: Statistical analyses.**
Complete results of statistical analyses for Figs. 3a-d, 5d-e, 6a-c, 6d, 7a,b,d, and Supplementary Fig. 8.
