## Supplementary Table 19 for "Translational profiling of mouse dopaminoceptive neurons reveals a role of PGE2 in dorsal striatum"

### Figure 3

| Figure | Variable | Group | n mice | n data points | Normality test | Mean | SEM | Statistical analysis | Comparison | | Test value | *P* value |
|---|---|---|---|---|---|---|---|---|---|---|---|---|
| 3a | Iso2a/Iso1 | DS | 5 | 5 | Small | 0,267 | 0,0174 | Mann Whitney's test two-tailed | DS vs NAc | | U = 0 | 0,008 |
| | | NAc | 5 | 5 | Small | 1,022 | 0,0798 | | | | | |
| 3b | Tbr1 | DS | 6 | 6 | Small | 100 | 21 | Mann Whitney's test two-tailed | PFC vs DS | | U = 0 | 0,002 |
| | | PFC | 6 | 6 | Small | 2111 | 79 | | | | | |
| | Ppp2r2b | DS | 6 | 6 | Small | 95 | 6 | Mann Whitney's test two-tailed | PFC vs DS | | U = 0 | 0,002 |
| | | PFC | 6 | 6 | Small | 221 | 8 | | | | | |
| | Shank2 | DS | 6 | 6 | Small | 100 | 4 | Mann Whitney's test two-tailed | PFC vs DS | | U = 0 | 0,002 |
| | | PFC | 6 | 6 | Small | 398 | 57 | | | | | |
| | Tac2 | DS | 6 | 6 | Small | 128 | 42 | Mann Whitney's test two-tailed | PFC vs DS | | U = 5 | 0,041 |
| | | PFC | 6 | 6 | Small | 291 | 30 | | | | | |
| 3c | Wfs1 | DS | 6 | 6 | Small | 99 | 5 | Mann Whitney's test two-tailed | DS vs NAc | | U = 0 | 0,002 |
| | | NAc | 6 | 6 | Small | 249 | 31 | | | | | |
| | Ahi1 | DS | 6 | 6 | Small | 120 | 20 | Mann Whitney's test two-tailed | DS vs NAc | | U = 0,5 | 0,004 |
| | | NAc | 6 | 6 | Small | 393 | 68 | | | | | |
| | Gda | DS | 6 | 6 | Small | 100 | 6 | Mann Whitney's test two-tailed | DS vs NAc | | U = 0 | 0,002 |
| | | NAc | 6 | 6 | Small | 214 | 29 | | | | | |
| | Dlk1 | DS | 6 | 6 | Small | 100 | 25 | Mann Whitney's test two-tailed | DS vs NAc | | U = 0 | 0,002 |
| | | NAc | 6 | 6 | Small | 2410 | 850 | | | | | |
| | Drd3 | DS | 4 | 4 | Small | 100 | 17 | Mann Whitney's test two-tailed | DS vs NAc | | U = 0 | 0,03 |
| | | NAc | 4 | 4 | Small | 985 | 278 | | | | | |
| | Arghap36 | DS | 4 | 4 | Small | 100 | 21 | Mann Whitney's test two-tailed | DS vs NAc | | U = 0 | 0,03 |
| | | NAc | 4 | 4 | Small | 2339 | 464 | | | | | |
| 3d | Cacng1 | DS | 6 | 6 | Small | 100 | 24 | Mann Whitney's test two-tailed | NAc vs DS | | U = 0 | 0,002 |
| | | NAc | 6 | 6 | Small | 570 | 134 | | | | | |
| | Hpca | DS | 6 | 6 | Small | 100 | 16 | Mann Whitney's test two-tailed | NAc vs DS | | U = 0 | 0,002 |
| | | NAc | 6 | 6 | Small | 258 | 23 | | | | | |
| | Atp2b1 | DS | 6 | 6 | Small | 100 | 16 | Mann Whitney's test two-tailed | NAc vs DS | | U = 0 | 0,002 |
| | | NAc | 6 | 6 | Small | 220 | 10 | | | | | |
| | Slc24a2 | DS | 6 | 6 | Small | 100 | 11 | Mann Whitney's test two-tailed | NAc vs DS | | U = 0 | 0,002 |
| | | NAc | 6 | 6 | Small | 225 | 9 | | | | | |
| | Pde10a | DS | 6 | 6 | Small | 100 | 18 | Mann Whitney's test two-tailed | NAc vs DS | | U = 0 | 0,002 |
| | | NAc | 6 | 6 | Small | 340 | 27 | | | | | |

### Figure 5

| Figure | Variable | Group | n mice | n data points | Normality test | Mean | SEM | Statistical analysis | Comparison | | Test value | *P* value |
|---|---|---|---|---|---|---|---|---|---|---|---|---|
| 5d | Ptger1 | NAc | 18 | 6 | Small | 1,01 | 0,06 | Mann Whitney's test two-tailed | NAc vs DS | | U = 1 | **0,007** |
| | | DS | 15 | 5 | Small | 1,33 | 0,07 | | | | | |
| | | NAc D1 | 6 | 6 | Small | 1,31 | 0,35 | Mann Whitney's test two-tailed | NAc vs DS | | U = 10 | **0,43** |
| | | DS D1 | 5 | 5 | Small | 2,65 | 1,17 | | | | | |
| | | NAc D2 | 6 | 6 | Small | 2,18 | 1,23 | Mann Whitney's test two-tailed | NAc vs DS | | U = 0 | **0,01** |
| | | DS D2 | 4 | 4 | Small | 68,78 | 18,49 | | | | | |
| 5e | Ptger2 | NAc | 18 | 6 | Small | 1,03 | 0,10 | Mann Whitney's test two-tailed | NAc vs DS | | U = 0 | **0,004** |
| | | DS | 15 | 5 | Small | 2,09 | 0,10 | | | | | |
| | | NAc D1 | 6 | 6 | Small | 0,58 | 0,52 | Mann Whitney's test two-tailed | NAc vs DS | | U = 3 | **0,02** |
| | | DS D1 | 5 | 5 | Small | 17,86 | 12,49 | | | | | |
| | | NAc D2 | 6 | 6 | Small | 0,80 | 0,52 | Mann Whitney's test two-tailed | NAc vs DS | | U = 0 | **0,01** |
| | | DS D2 | 4 | 4 | Small | 27,72 | 7,45 | | | | | |

### Figure 6a-c

| Figure | Variable | Group | n mice | n data points | | Mean | | Statistical analysis | Comparison | DF | Test value | *P* value |
|---|---|---|---|---|---|---|---|---|---|---|---|---|
| 6a | pPKA substrates | Vehicle ip | 5 | 5 | Small | 100,00 | 2,26 | Mann Whitney's test two-tailed | NAc vs DS | | U = 2 | 0,01 |
| | | Misoprostol ip | 7 | 7 | Small | 139,90 | 14,54 | | | | | |
| 6b | Rotarod 24 rpm latency to fall | Vehicle (DS) | 9 | 9 | Yes | 13,11 | 1,34 | t test two-tailed | Vehicle vs misoprostol | 14 | t = 2,209 | 0,04 |
| | | Misoprostol (DS) | 7 | 7 | Yes | 9,36 | 0,84 | | | | | |
| 6c | Y maze learning phase Correct choice (%) | Vehicle (osmopump ip) | 20 | 120 | | | | Two-way ANOVA repeated measures | Interaction | 5 ; 210 | F = 1,25 | 0,29 |
| | | Misoprostol (osmopump ip) | 24 | 144 | | | | | Time | 5 ; 210 | F = 8,47 | < 0.0001 |
| | | | | | | | | | Treatment | 1 ; 42 | F = 0,69 | 0,41 |
| | | | | | | | | | Subjects (matching) | 42 ; 210 | F = 2,57 | < 0.0001 |
| | | Trial 1 Vehicle pump ip | 20 | 20 | | 54,00 | | Holm Sidak's multiple comparisons test | Misoprostol vs Vehicle | 252 | t = 1,81 | 0,36 |
| | | Trial 1 Misoprostol pump ip | 24 | 24 | | 64,17 | | | | | | |
| | | Trial 2 Vehicle pump ip | 20 | 20 | | 64,00 | | | Misoprostol vs Vehicle | 252 | t = 0,33 | 1,00 |
| | | Trial 2 Misoprostol pump ip | 24 | 24 | | 65,83 | | | | | | |
| | | Trial 3 Vehicle pump ip | 20 | 20 | | 62,50 | | | Misoprostol vs Vehicle | 252 | t = 1,48 | 0,60 |
| | | Trial 3 Misoprostol pump ip | 24 | 24 | | 70,83 | | | | | | |
| | | Trial 4 Vehicle pump ip | 20 | 20 | | 72,00 | | | Misoprostol vs Vehicle | 252 | t = 0,24 | > 0,99 |
| | | Trial 4 Misoprostol pump ip | 24 | 24 | | 73,33 | | | | | | |
| | | Trial 5 Vehicle pump ip | 20 | 20 | | 78,00 | | | Misoprostol vs Vehicle | 252 | t = 0,09 | > 0,99 |
| | | Trial 5 Misoprostol pump ip | 24 | 24 | | 77,50 | | | | | | |
| | | Trial 6 Vehicle pump ip | 20 | 20 | | 79,00 | | | Misoprostol vs Vehicle | 252 | t = 0,86 | 0,95 |
| | | Trial 6 Misoprostol pump ip | 24 | 24 | | 74,17 | | | | | | |
| | Y maze reversal phase Correct choice (%) | Vehicle (pump ip) | 20 | 80 | | | | Two-way ANOVA repeated measures | Interaction | 3 ; 123 | F = 1,77 | 0,16 |
| | | Misoprostol (pump ip) | 23 | 92 | | | | | Time | 4 ; 123 | F = 33,97 | < 0.0001 |
| | | | | | | | | | Treatment | 1 ; 41 | F = 17,20 | 0.0002 |
| | | | | | | | | | Subjects (matching) | 41 ; 123 | F = 1,50 | 0,046 |
| | | Trial 9 Vehicle pump ip | 20 | 20 | | 44,00 | | Holm Sidak's multiple comparisons test | Misoprostol vs Vehicle | 164 | t = 3,53 | 0,002 |
| | | Trial 9 Misoprostol pump ip | 23 | 23 | | 60,00 | | | | | | |
| | | Trial 10 Vehicle pump ip | 20 | 20 | | 57,00 | | | Misoprostol vs Vehicle | 164 | t = 2,58 | 0,042 |
| | | Trial 10 Misoprostol pump ip | 23 | 23 | | 68,70 | | | | | | |
| | | Trial 11 Vehicle pump ip | 20 | 20 | | 72,00 | | | Misoprostol vs Vehicle | 164 | t = 0,61 | 0,955 |
| | | Trial 11 Misoprostol pump ip | 23 | 23 | | 74,78 | | | | | | |
| | | Trial 12 Vehicle pump ip | 20 | 20 | | 74,00 | | | Misoprostol vs Vehicle | 164 | t = 2,86 | 0,019 |
| | | Trial 12 Misoprostol pump ip | 23 | 23 | | 86,96 | | | | | | |

### Figure 6d

| Figure | Variable | Group | n | n data | Mean | Statistical analysis | Comparison | DF | Test value | *P* value |
|---|---|---|---|---|---|---|---|---|---|---|
| 6d | Y maze learning phase Correct choice (%) | Vehicle (osmopump DS) | 10 | 80 | | Two-way ANOVA repeated measures | Interaction | 7 ; 119 | F = 1,52 | 0,16 |
| | | Misoprostol (osmopump DS) | 9 | 72 | | | Time | 7 ; 119 | F = 8,57 | **< 0.0001** |
| | | | | | | | Treatment | 1 ; 17 | F = 12,17 | **0.003** |
| | | | | | | | Subjects (matching) | 17 ; 119 | F = 1,2 | 0,28 |
| | | Trial 1 Vehicle DS | 10 | 10 | 56,00 | Holm Sidak's multiple comparisons test | Misoprostol vs Vehicle | 136 | t = 0,97 | 0,80 |
| | | Trial 1 Misoprostol DS | 9 | 9 | 62,22 | | | | | |
| | | Trial 2 Vehicle DS | 10 | 10 | 56,00 | | Misoprostol vs Vehicle | 136 | t = 0,069 | 1,00 |
| | | Trial 2 Misoprostol DS | 9 | 9 | 55,56 | | | | | |
| | | Trial 3 Vehicle DS | 10 | 10 | 58,00 | | Misoprostol vs Vehicle | 136 | t = 0,035 | 1,00 |
| | | Trial 3 Misoprostol DS | 9 | 9 | 57,78 | | | | | |
| | | Trial 4 Vehicle DS | 10 | 10 | 60,00 | | Misoprostol vs Vehicle | 136 | t = 1,39 | 0,60 |
| | | Trial 4 Misoprostol DS | 9 | 9 | 68,89 | | | | | |
| | | Trial 5 Vehicle DS | 10 | 10 | 66,00 | | Misoprostol vs Vehicle | 136 | t = 0,45 | 0,96 |
| | | Trial 5 Misoprostol DS | 9 | 9 | 68,89 | | | | | |
| | | Trial 6 Vehicle DS | 10 | 10 | 58,00 | | Misoprostol vs Vehicle | 136 | t = 2,39 | 0,12 |
| | | Trial 6 Misoprostol DS | 9 | 9 | 73,33 | | | | | |
| | | Trial 7 Vehicle DS | 10 | 10 | 64,00 | | Misoprostol vs Vehicle | 136 | t = 3,19 | **0,01** |
| | | Trial 7 Misoprostol DS | 9 | 9 | 84,44 | | | | | |
| | | Trial 8 Vehicle DS | 10 | 10 | 76,00 | | Misoprostol vs Vehicle | 136 | t = 2,36 | 0,12 |
| | | Trial 8 Misoprostol DS | 9 | 9 | 91,11 | | | | | |
| | Y maze reversal phase Correct choice (%) | Vehicle (osmopump DS) | 10 | 40 | | Two-way ANOVA repeated measures | Interaction | 7 ; 119 | F = 1,51 | 0,22 |
| | | Misoprostol (osmopump DS) | 9 | 36 | | | Time | 7 ; 119 | F = 19,06 | **< 0.0001** |
| | | | | | | | Treatment | 1 ; 17 | F = 13,62 | **0.002** |
| | | | | | | | Subjects (matching) | 17 ; 119 | F = 1,76 | 0,06 |
| | | Trial 9 Vehicle DS | 10 | 10 | 36,0 | Holm Sidak's multiple comparisons test | Misoprostol vs Vehicle | 68 | t = 2,37 | 0,061 |
| | | Trial 9 Misoprostol DS | 9 | 9 | 57,8 | | | | | |
| | | Trial 10 Vehicle DS | 10 | 10 | 62,0 | | Misoprostol vs Vehicle | 68 | t = 2,20 | 0,062 |
| | | Trial 10 Misoprostol DS | 9 | 9 | 82,2 | | | | | |
| | | Trial 11 Vehicle DS | 10 | 10 | 56,0 | | Misoprostol vs Vehicle | 68 | t = 3,58 | **0,003** |
| | | Trial 11 Misoprostol DS | 9 | 9 | 88,9 | | | | | |
| | | Trial 12 Vehicle DS | 10 | 10 | 88,0 | | Misoprostol vs Vehicle | 68 | t = 0,82 | 0,41 |
| | | Trial 12 Misoprostol DS | 9 | 9 | 95,6 | | | | | |

### Figure 7

| Figure | Variable | Group | n mice | n data points | Normality test | Mean | SEM | Statistical analysis | Comparison | DF | Test value | *P* value |
|---|---|---|---|---|---|---|---|---|---|---|---|---|
| 7a,b | AUC Fiber photometry (new environment minus baseline) | Vehicle (i.p.) | 6 | 6 | Small | | | One-sample Wilcoxon two-tailed | New environment vs baseline | | | **0,03** |
| | | Misoprostol (i.p.) | 6 | 6 | Small | | | | New environment vs baseline | | | **0,16** |
| 7d | Haloperidol-induced catalepsy latency to fall (sec) | Vehicle (ip) | 9 | 81 | | | | Two-way ANOVA repeated measures | Interaction | 8 ; 128 | F = 9,75 | **< 0.0001** |
| | | Misoprostol (ip) | 9 | 81 | | | | | Time | 8 ; 128 | F = 18,17 | **< 0.0001** |
| | | | | | | | | | Treatment | 1 ; 16 | F = 34,53 | **< 0.0001** |
| | | | | | | | | | Subjects (matching) | 16 ; 128 | F = 3,68 | **< 0.0001** |
| | | Trial 1 Vehicle | 10 | 10 | | 9,56 | | Holm Sidak's multiple comparisons test | Misoprostol vs Vehicle | 144 | t = 1,5 | 0,99 |
| | | Trial 1 Misoprostol | 9 | 9 | | 7,22 | | | | | | |
| | | Trial 2 Vehicle | 10 | 10 | | 16,89 | | | Misoprostol vs Vehicle | 144 | t = 0,078 | 0,99 |
| | | Trial 2 Misoprostol | 9 | 9 | | 18,11 | | | | | | |
| | | Trial 3 Vehicle | 10 | 10 | | 40,89 | | | Misoprostol vs Vehicle | 144 | t = 0,97 | 0,70 |
| | | Trial 3 Misoprostol | 9 | 9 | | 25,78 | | | | | | |
| | | Trial 4 Vehicle | 10 | 10 | | 47,89 | | | Misoprostol vs Vehicle | 144 | t = 1,30 | 0,58 |
| | | Trial 4 Misoprostol | 9 | 9 | | 27,67 | | | | | | |
| | | Trial 5 Vehicle | 10 | 10 | | 83,11 | | | Misoprostol vs Vehicle | 144 | t = 4,02 | **0,0005** |
| | | Trial 5 Misoprostol | 9 | 9 | | 20,44 | | | | | | |
| | | Trial 6 Vehicle | 10 | 10 | | 110,30 | | | Misoprostol vs Vehicle | 144 | t = 5,45 | **< 0.0001** |
| | | Trial 6 Misoprostol | 9 | 9 | | 25,44 | | | | | | |
| | | Trial 7 Vehicle | 10 | 10 | | 125,30 | | | Misoprostol vs Vehicle | 144 | t = 6,52 | **< 0.0001** |
| | | Trial 7 Misoprostol | 9 | 9 | | 23,67 | | | | | | |
| | | Trial 8 Vehicle | 10 | 10 | | 143,20 | | | Misoprostol vs Vehicle | 144 | t = 6,79 | **< 0.0001** |
| | | Trial 8 Misoprostol | 9 | 9 | | 37,33 | | | | | | |
| | | Trial 9 Vehicle | 10 | 10 | | 118,20 | | | Misoprostol vs Vehicle | 144 | t = 4,57 | **< 0.0001** |
| | | Trial 9 Misoprostol | 9 | 9 | | 47,00 | | | | | | |

### Supplementary Fig. 8

| Figure | Variable | Group | n | n data | | Statistical analysis | Comparison | DF | Test value | *P* value |
|---|---|---|---|---|---|---|---|---|---|---|
| **Sup. Fig. 8a** | Rotarod training latency to fall | Vehicle (osmopump ip) | 14 | 300 | | Two-way ANOVA repeated measures | Interaction | 13 ; 562 | F = 0,57 | 0,88 |
| | | Misoprostol (osmopump ip) | 14 | 290 | | | Time | 13 ; 562 | F = 31,56 | **< 0.0001** |
| | | | | | | | Treatment | 1 ; 562 | F = 0,68 | 0,41 |
| **Sup. Fig. 8b** | Rotarod training latency to fall | Vehicle (osmopump DS) | 10 | 140 | | Two-way ANOVA repeated measures | Interaction | 13 ; 221 | F = 2,34 | 0,006 |
| | | Misoprostol (osmopump DS) | 9 | 126 | | | Time | 13 ; 221 | F = 28,68 | **< 0.0001** |
| | | | | | | | Treatment | 1 ; 17 | F = 0,64 | 0,43 |
